# Simulation-free approximate Bayesian computation for stochastic reaction networks

**DOI:** 10.64898/2026.10.19.760104

**Authors:** Zekai Li, Mauricio Barahona, Philipp Thomas

## Abstract

Uncovering stochastic reaction networks from data is central to single-cell biology. Simulation-based inference has evolved into a flexible paradigm for this purpose, circumventing the computational intractability of the likelihood. Notably, approximate Bayesian computation performs this inference by carrying out extensive Gillespie simulations, usually at a high computational cost. Here, we propose a simulation-free approximate Bayesian computation (SFABC) framework that avoids both simulation and explicit likelihood solutions of the network dynamics. Instead, SFABC exploits the Chemical Master Equation to formulate a tractable convex optimization with ensuing constraints, which are then used to accept sampled parameters consistent with the observed data. The resulting posterior approximation comes with theoretical guarantees: it can be tightened systematically, and it provably converges to the exact Bayesian posterior distributions. Using synthetic benchmarks, we show that SFABC matches the accuracy of standard simulation-based algorithms at a substantially reduced computational cost. We then showcase the applicability of SFABC to experimental systems biology by analyzing gene transcription bursting kinetics from snapshot, time-resolved, and cell-cycle-resolved single-cell data.

## I. INTRODUCTION

With the advent of single-cell sequencing technologies, large-scale transcriptomic datasets have become available, enabling the study of single-cell dynamics and providing insights into complex biological mechanisms spanning gene expression heterogeneity^1–6^, molecular interaction networks^7–11^, or differentiation trajectories^12–14^. To capture the stochasticity arising from a low number of molecules per cell, these dynamics are often modeled as stochastic reaction networks (SRNs) that can be analyzed using the Chemical Master Equation (CME) and Gillespie’s stochastic simulation algorithm^15^, in which reaction events occur randomly with propensities determined by the network structure and reaction rates^16–20^. Inferring these reaction rate parameters is therefore crucial for understanding the underlying biological mechanisms, but it remains challenging given the stochastic nature of cellular dynamics^21,22^.

Numerous methods for parameter inference from single-cell data have been developed over the past decades. Likelihood-based approaches estimate parameters by explicitly modeling the probability of the observed data through maximum likelihood estimation^2,6,23^ or Bayesian inference^24–26^. However, likelihood evaluation requires analytical or numerical solutions to the CME that present an infinite system of equations and can become intractable even for relatively simple networks^21^ and require further approximations^27–29^. Likelihood-free methods offer a tangible solution by avoiding explicit evaluation of the likelihood. Although moment approximations provide a frequentist likelihood-free route^27,30,31^, most Bayesian likelihood-free approaches still rely on stochastic simulations to generate realizations of stochastic reaction networks^32–37^, making simulation-based inference (SBI) a central paradigm in likelihood-free Bayesian inference^38^.

A limitation of SBI is the trade-off between estimation accuracy and computational cost, since improving accuracy typically requires more simulations. As a classical SBI approach, approximate Bayesian computation (ABC) estimates the posterior by repeatedly simulating data under parameter proposals and comparing the simulated data with observations^39–43^. Although ABC has provable convergence to the true posterior, this convergence comes at the cost of rapidly increasing computational burden^44–46^, despite the development of more efficient sampling schemes and other algorithmic improvements^47–53^. Alternatively, recent advances in machine learning have accelerated SBI through both surrogate models of the simulation process^54–61^ and neural posterior estimation methods that learn the posterior directly from simulated data^35,62–64^. However, these neural approximations can introduce inductive bias that forfeits convergence guarantees, and so the quality of SBI must be assessed on a case-by-case basis since it critically depends on the number of simulations available for training^65^.

In this work, we introduce simulation-free approximate Bayesian computation (SFABC) for SRNs, a novel framework for likelihood-free Bayesian inference that circumvents the need for stochastic simulations. Grounded in the CME, the system of coupled differential equations that governs the dynamics of an SRN, SFABC checks whether a sampled parameter set is compatible with a candidate state probability obtained from the observed data (Fig. 1). We prove that the approximated posterior from SFABC converges to the true posterior, subject to several limits that involve zero tolerance. Using synthetic datasets, we demonstrate that SFABC is compatible with various sampling schemes and achieves accurate posterior estimates, while substantially reducing computational cost as the number of observed cells increases. Applied to snapshot scRNA-seq data, SFABC uncovers differences in transcriptional kinetics between genes with different promoter architectures and reveals their posterior dependence structure. With temporal data, SFABC enables inference of kinetic rates on an absolute time scale and recovers associations across data modalities. Finally, we apply SFABC to distinguish among competing models of cell cycle-dependent regulation.

**FIG. 1.**
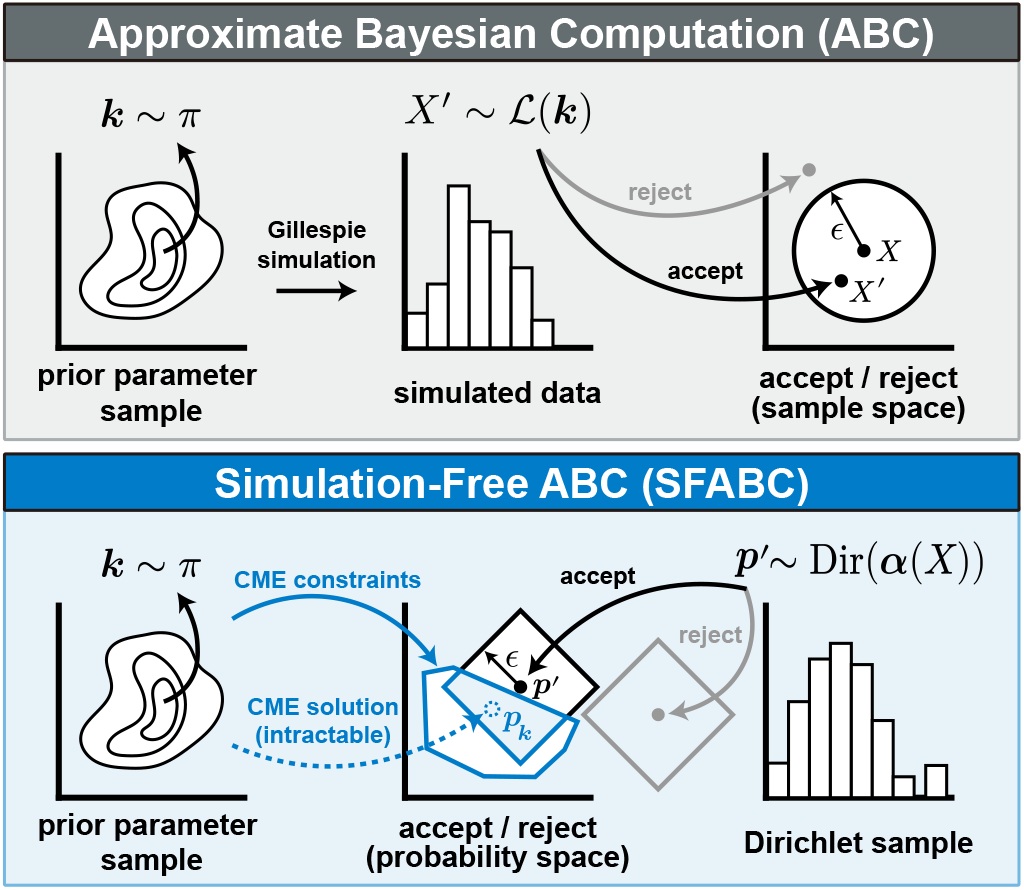
Comparison of ABC and SFABC for SRNs. ABC simulates data *X*^*′*^ from the CME likelihood *ℒ* using the Gillespie algorithm given a parameter set ***k*** drawn from the prior *π*. Parameters producing simulations *X*^*′*^ close (*ϵ*-neighborhood) to the observed data *X* are accepted to approximate the posterior distribution (*SI Text 1*). SFABC draws probability distributions ***p***^*′*^ using Dirichlet resampling of the data. A parameter set ***k*** is accepted if the *ϵ*-neighborhood of the candidate probability ***p***^*′*^ contains probability vectors that satisfy the CME constraints, which can be checked by convex optimization. The CME constraints present a relaxation containing the intractable CME solution ***p***_***k***_ that can be systematically refined.

## II. RESULTS

We consider Bayesian inference of rate parameters 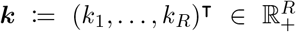 in an SRN with molecular species ***Y*** := (*Y*_*s*_)_*s*∈*S*′_, where *S*′ is the species index set, participating in *r* = 1, …, *R* reactions

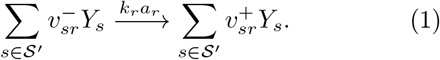

In reaction *r*, 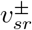 molecules of species *Y*_*s*_ are consumed and produced, respectively, and *a*_*r*_ is the state-dependent, unit-rate propensity function. Let us consider a dataset *X* := (***x***_1_, …, ***x***_*N*_) ℕ^|*S*|*×N*^ that contains *N* independent observations of counts of the observable species *S* ⊆ *S*′ in cells at time points 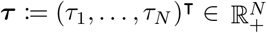. Then the posterior distribution of the rate parameters ***k*** is:

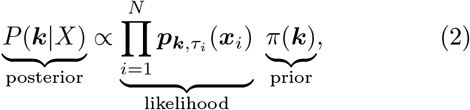

where ***p***_***k***,*t*_(***x***) denotes the marginalized probability over the observed species, and the full state probability ***p***_***k***,*t*_ satisfies the CME

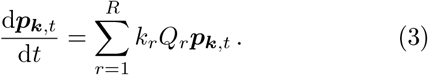

Here, the unit-rate transition matrix is defined by *Q*_*r*_(***x, y***) := *a*_*r*_(***y***)(*I*_***y***+***v***_ (***x***) *− I*_***y***_(***x***)), where 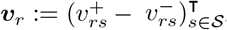 is the stoichiometric vector and ***x, y*** ∈ ***X*** with ***X*** ⊆ ℕ^|*S′*|^ denoting the full state space. In the following, we assume that the state space is irreducible; hence, there exists a unique solution to the CME^66^.

Calculating the explicit state probability ***p***_***k***,*t*_, and hence the likelihood, requires solving the CME in (3), which is often mathematically demanding because the underlying state space is typically infinite, giving rise to an infinite system of coupled differential equations. While likelihood-based MCMC is applicable only to the limited class of SRNs with tractable likelihoods, simulation-based methods such as ABC avoid explicit likelihood evaluation by using stochastic simulations. However, repeated simulations can become computationally prohibitive. We therefore introduce a simulation-free ABC framework that uses CME-derived constraints to directly assess the compatibility of candidate parameters with the observed data (see Fig. 1), requiring neither stochastic simulation nor explicit likelihood evaluation.

### A. SFABC Provides Likelihood- and Simulation-Free Posterior Estimation

The core idea of SFABC is to accept (or reject) a prior parameter sample ***k*** based on whether the resampled data is compatible (or incompatible) with the solution of the CME. A computational challenge with this approach is that the CME typically evolves over an unbounded state space larger than the one observed in the data. We circumvent this issue by formulating a tractable convex linear optimization problem that can be solved using standard solvers^67,68^. Specifically, we consider only a finite underdetermined set of equations in (3) involving probabilities over a subset ***M*** ⊆ ***X*** of the state space. To deal with the time-derivatives involved in transient problems, we integrate (3) against a range of suitable time-dependent test functions which provides constraints on a finite time grid involving the observation times (see *SI Text 2*). The resulting relaxation approximation represents a set 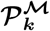 of plausible CME solutions given a finite set of equations that also contains the true solution and can be tightened by increasing the number of equations through ***M***.

Let 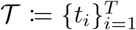 denote the set of unique observation times in ***τ***. For each *t*, let *B*_*t*_ be the number of unique observed states at time *t* with 1 *≤ B*_*t*_ *≤ N*_*t*_, and we denote the unique states by 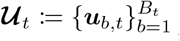 with numbers of occurrence *n*_1,*t*_, …, *n*_*B, t*_ satisfying 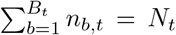. The SFABC rejection sampling algorithm is then defined as follows:

#### SFABC rejection sampling (SFABC-Rej)

*S1* Sample a parameter set ***k*** from the prior *π*.

*S2* Sample probability distributions 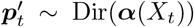 with 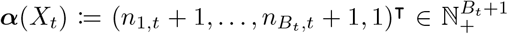 for each observation time *t* ∈ *T*.

*S3* Test if there exists a CME relaxation 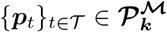 that satisfies max 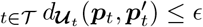. If such a solution exists, accept ***k***. Otherwise, go to *S1*.

*S4* Weigh the accepted sample 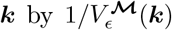 where the volume of acceptance region 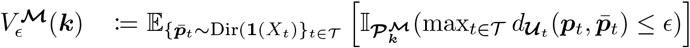, where 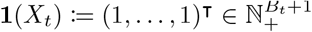.

Here, *S1* samples a parameter ***k*** from the prior. In *S2*, we sample a probability distribution from a Dirichlet distribution whose first *B*_*t*_ components correspond to the probabilities over the observed states ***U***_*t*_, and the last component aggregates the total probability mass over all unobserved states. This step captures finite-sample variability. Step *S3*, called the feasibility test, tests whether the probability distributions 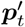 for all *t* ∈ *T* are close (within an *ϵ* neighborhood) to a CME relaxation with parameters ***k***. The distance 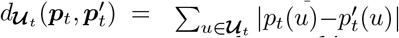 is the 1-norm computed over the observed states ***U***_*t*_ or a suitably chosen distance of summary statistics (*SI Text 4*). Finally, *S4* weighs the accepted parameters by the inverse volume of the acceptance region obtained by repeated sampling from uniform Dirichlet probability distributions (see *Materials and Methods*). A generalization of this algorithm is Sequential Monte Carlo (SFABC-SMC) that can improve sampling efficiency for parameter inference and model selection (*SI Text 3*).

The construction is a computationally tractable linear program relaxation of the CME solution that converges to it as the subset of equations covers the whole probability distribution and temporal dimension, namely as ***M*** → ***X*** and the time discretization decreases to zero. The accepted samples in SFABC-Rej therefore can, in principle, approximate the true posterior (2) arbitrarily closely as *ϵ* → 0 (see convergence proof in *Materials and Methods*). In practice, the resulting posterior approximations are expected to be wider than the limiting true posteriors since the relaxation approximation contains the true CME solution. In the following, we validate that SFABC provides accurate posterior approximations and demonstrate how it can be adapted to a broad range of single-cell transcriptomic data.

### B. SFABC Accurately and Robustly Estimates the Posterior Distribution with Lower Computational Cost

First, we validate the accuracy of SFABC’s posterior estimate against exact posterior samples obtained using a likelihood-based MCMC sampler (see *SI Text 4*) and we benchmark its performance against standard simulation-based ABC. We consider synthetic snapshot data from the telegraph model (see *SI Text 5* and illustration in Fig. 2A). At steady state, the time derivative in (3) is zero, which reduces the CME to algebraic constraints:

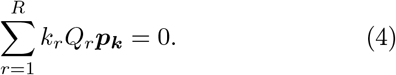

**FIG. 2.**
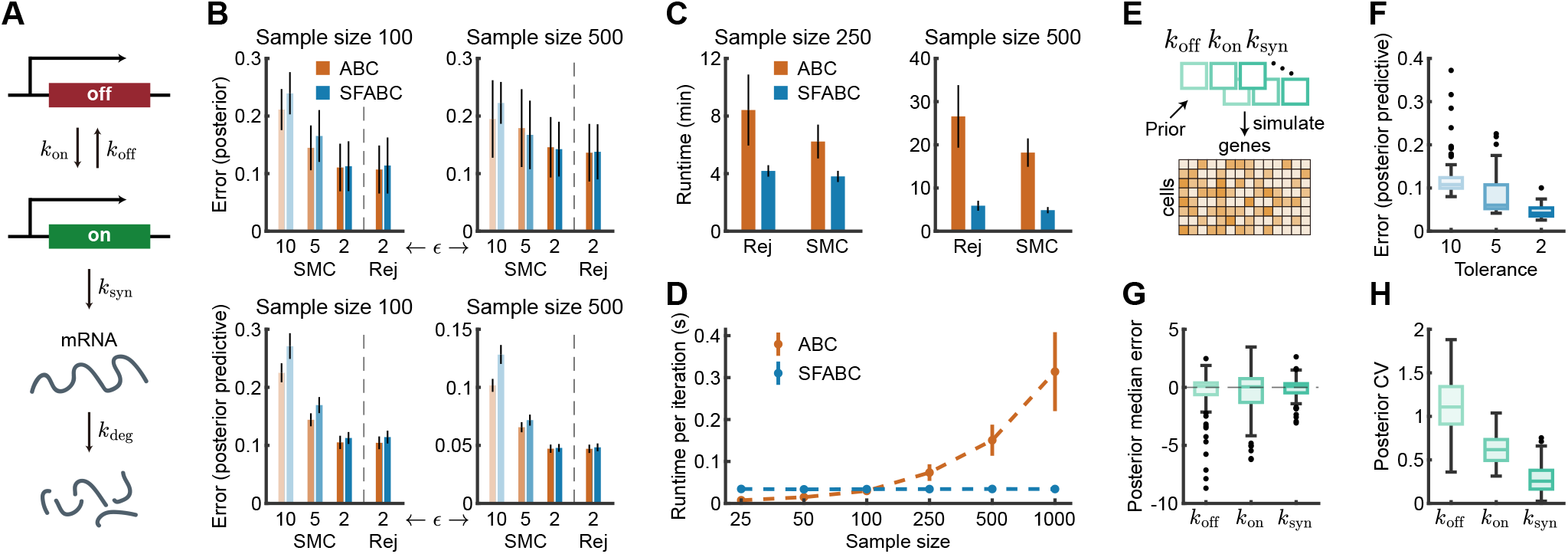
Validation of SFABC with Steady-State Synthetic Data from the Telegraph Model. **(A)** Illustration of the telegraph model. **(B)** Validation of SFABC algorithms with sample sizes 100 and 500 drawn from the analytical steady-state distribution^69^ (*k*_off_ = 2, *k*_on_ = 8, *k*_syn_ = 5, *k*_deg_ = 1). For each sample size, 20 datasets are simulated, and 500 posterior samples are collected per dataset using MCMC, ABC-Rej, ABC-SMC, SFABC-Rej, and SFABC-SMC sampling algorithms. The SMC algorithms use tolerance sequence *ϵ* = {10, 5, 2} and the rejection samplers use *ϵ* = 2, and other inference setups are provided in S1 - S4 in *SI Table S1*. Top panels: mean Kolmogorov–Smirnov (KS) statistic between the marginal distributions of SFABC posterior samples and MCMC samples. Bottom panels: mean KS statistic between each observed dataset and 500 synthetic datasets drawn from the corresponding posterior predictive distribution. Bars indicate mean metric ± standard deviation. **(C)** Total wall time for sample sizes 250 and 500. Experiments are repeated over 20 datasets, and bars indicate the mean time ± standard deviation. **(D)** Wall time per iteration (simulation for ABC and feasibility test for SFABC) with sample sizes from 25 to 1,000. Points and bars indicate mean wall time ± standard deviation over 100 iterations. **(E)** Validation of SFABC inference across parameter sets. Parameters *k*_off_, *k*_on_, *k*_syn_ are drawn independently from 10^Uniform(*−*1,1)^ and used to simulate datasets of size 1,000 from the steady-state distribution. We retain only datasets with mean expression greater than 0.5, yielding 100 parameter sets and synthetic datasets. For each dataset, we use SFABC-SMC to collect 100 posterior samples with tolerance sequence *ϵ* = {10, 5, 2} (full setup in S5 in *SI Table S1*). **(F)** KS statistic between observed data and 100 synthetic datasets drawn from the posterior predictive distribution. **(G)** Difference between the true parameter and posterior median with *ϵ* = 2. **(H)** Coefficient of variation (CV) of marginal posterior distributions with *ϵ* = 2.

Restricting the above to a finite subset of equations involving the observed state probabilities and the distance to the sampled histogram then provides constraints that represent our relaxation approximation (*SI Text 2*).

We find that SFABC accurately estimates the posterior distribution across datasets with varying sample sizes (Fig. 2B). The approximation errors of SFABC-SMC decrease with tolerance and achieve accuracy similar to that of SFABC-Rej, consistent with SFABC’s guarantee of convergence to the true posterior (blue bars). This is quantified by the Kolmogorov–Smirnov (KS) distance between the exact posterior obtained by MCMC and the SFABC approximation (top), and similarly, the KS distance between the snapshot data distribution and SFABC’s posterior predictive distribution (bottom row). Posterior accuracy improves with the number of equations used in the optimization and as the tolerance decreases, but this also decreases the acceptance probability as expected (***SI Fig. S1***). This accuracy is comparable to that of the standard ABC (Fig. 2B, red bars), despite fundamental differences in the rejection step, and is independently verified for our SMC implementations, indicating that the relaxation employed by SFABC (*SI Text 4*) provides an excellent approximation to the CME.

While having similar accuracy, SFABC-Rej achieves a marked 50% reduction in computation time for sample size 250 and a 78% reduction for sample size 500 compared to ABC-Rej, and a similar scaling advantage is observed with the corresponding SMC samplers (Fig. 2C). The reduction in computational cost stems from the fact that the feasibility test with an LP solver at each SFABC iteration is virtually independent of sample size, in contrast to ABC’s simulation step (Fig. 2D).

Next, we assess the robustness of SFABC-SMC across a wide range of parameter settings. Across synthetic datasets obtained from 100 parameter sets sampled independently from an uninformative prior distribution (Fig. 2E, ***SI Fig. S2*** A), SFABC produces accurate posterior predictive distributions and unbiased parameter estimates, with improved agreement as the tolerance decreases (Fig. 2F-G). We observe that the pairwise posterior distributions from SFABC closely match those from ABC across datasets (***SI Fig. S2*** B-D). Despite identical priors, the relative posterior uncertainty is significantly lower for *k*_syn_ than for *k*_on_ and *k*_off_ (Fig. 2H). This is consistent with the intuition that *k*_syn_ sets expression levels^35^. Our results demonstrate the correctness, performance, and robust inference of SFABC.

### C. SFABC Enables Bayesian Analysis of Genome-wide Burst Kinetics

Next, we apply SFABC to analyze two scRNA-seq datasets in mouse fibroblast cells that differ substantially in sample size (224 cells in D1^6^; 8,916 in D2^70^; *SI Text 6*), and infer the underlying burst kinetics using the telegraph model. SFABC-SMC is calibrated with the tolerance sequences (setups S7 and S8 in *SI Table S1*) selected based on posterior predictive checks on a subset of genes (***SI Fig. S3***).

We first examine how the kinetic parameters depend on the promoter architecture. As reported previously^6^, genes with a TATA-binding element in their core promoter exhibit significantly higher burst sizes (BS = *k*_syn_*/k*_off_) than non-TATA-box genes, and are therefore considered noisier. We observe the same pattern in the SFABC posterior means for D1 (*P* = 1.73.01 × 10^*−*4^) and D2 (*P* = 9.58 × 10^*−*11^) (Fig. 3A). While the burst frequency (BF = *k*_on_*/k*_deg_ with *k*_deg_ ≡ 1) shows no difference in both datasets (Fig. 3C), several differences emerge in D2 that are not statistically significant in D1 due to low number of cells. These include lower gene activity (defined as GA = *k*_on_*/*(*k*_on_ + *k*_off_)) for TATA-box genes (*P* = 1.69 × 10^*−*2^) (Fig. 3B), and higher BS are reflected by difference between promoter types in D2 such a higher *k*_syn_ (*P* = 6.29 × 10^*−*6^) and lower *k*_off_ (*P* = 8.60 × 10^*−*3^) for TATA-box genes, while no difference in individual rates is observed in D1 (Fig. 3D-E). These findings suggest an additional role for TATA-containing promoters in modulating gene activity via promoter switching and thereby shaping overall burst kinetics distributions (Fig. 3F).

**FIG. 3.**
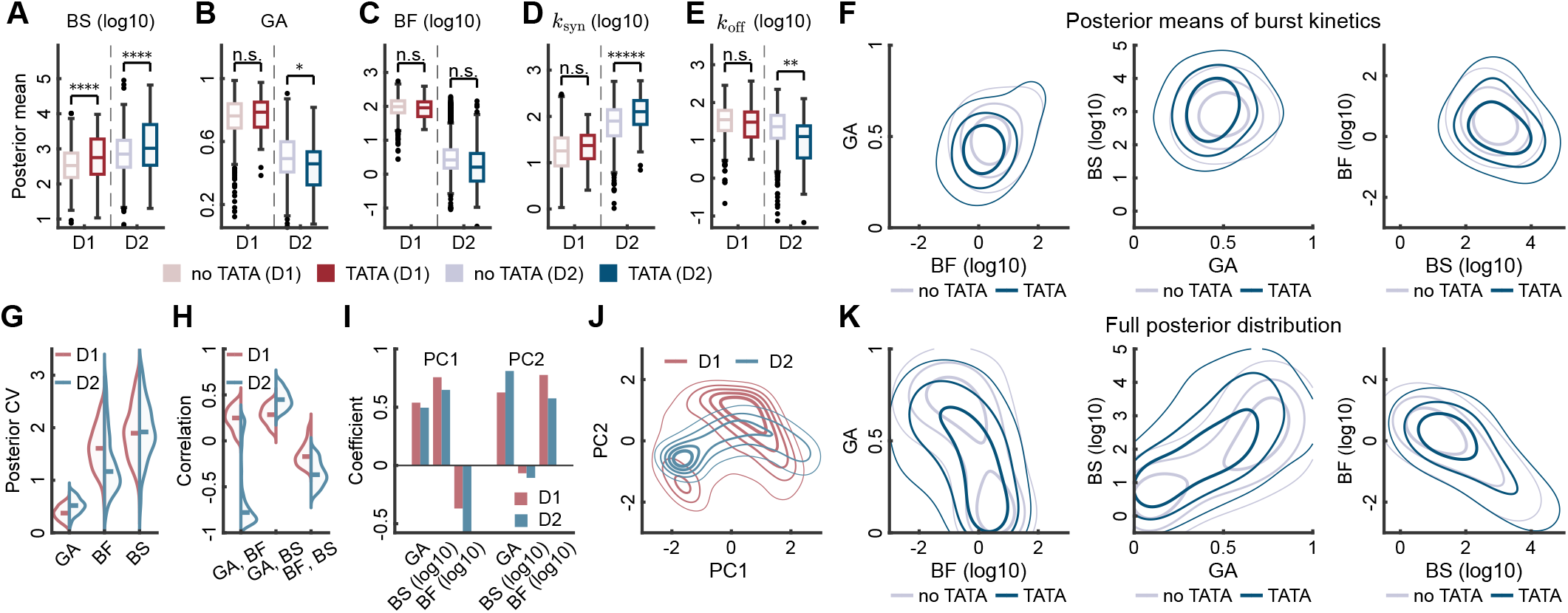
Inference of Burst Dynamics using the Telegraph Model from Snapshot scRNA-seq Data. **(A)-(E)** Posterior means of burst kinetics and individual rate parameters of the telegraph model (see cartoon in Fig. 2A), grouped by mouse fibroblast data (D1^6^, D2^70^) and promoter type (with or without a TATA element). **(F)** Smoothed contour plots of posterior means of gene activity (GA = *k*_on_*/*(*k*_on_ + *k*_off_)), burst size (BS = *k*_syn_*/k*_off_), and burst frequency (BF = *k*_on_*/k*_deg_) across genes (100 posterior samples per gene) in dataset D2, restricted to the 2.5th-97.5th percentiles. **(G)-(H)** CVs of marginal posterior distributions and pairwise correlations in posterior samples across genes in each dataset, restricted to the 10th-90th percentiles. **(I)** Coefficients of principle components. For each gene, principal component analysis is performed on the (across-gene) posterior samples of GA, log_10_(BS), and log_10_(BF) for each of D1 and D2. **(J)** Distribution of posterior samples projected to the plane of the first principal component (PC1) and the second one (PC2). **(K)** Smoothed contour plots of the full posterior distribution of GA, BF, and BS across genes in dataset D2, restricted to the 2.5th-97.5th percentiles. In all panels, SFABC-SMC is applied to 979 filtered genes (*SI Text 6*).

Although D2 contains many more cells than D1, the posterior uncertainty of the burst kinetics BS, GA, and BF does not decrease substantially (Fig. 3G). These measures could partly reflect differences between allele-specific expression in D1 and gene-level, effectively biallelic expression in D2 (*SI Text 6*), rather than posterior uncertainty alone. To investigate this, we first examine gene-level posterior dependence and find that D2 yields much stronger pairwise correlations (Fig. 3H). To assess whether this reflects a broader global structure in parameter identifiability, we next analyze the genome-wide posterior distribution that samples parameters across all genes. Principal component analysis of the burst kinetics (coefficients in Fig. 3I) shows that the first principal component explains more than 70% of the variance in D2, compared with only about 50% in D1 (Fig. 3J). The same pattern of posterior uncertainty was observed when D2 was subsampled to match the cell number in D1 (***SI Fig. S4***), suggesting that the first principal component quantifies uncertainty due to parameter identifiability, whereas the second component quantifies uncertainty due to finite sample size. Thus, sequencing more cells does not reduce posterior uncertainty but instead confines the posterior more strongly to a lower-dimensional manifold of burst kinetics, a behavior characteristic of the sloppiness in parameter identifiability^71^.

The sloppiness of burst kinetics is supported by comparing gene-level posterior means with the full posterior distributions that aggregate posterior samples across all genes. When considering posterior means (Fig. 3F), GA shows positive associations with BF and BS in both TATA-box and non-TATA-box genes (Pearson correlation coefficients 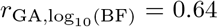 and 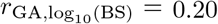 for TATA-box genes, and 0.53 and 0.15 for non-TATA-box genes). In contrast, BF and BS exhibit a weak inverse relationship 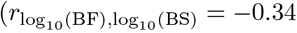 for TATA-box genes and − 0.37 for non-TATA-box genes), suggesting coordinated regulation of bursting kinetics at the between-gene level. In contrast, analysis of all posterior samples across genes reveals a stronger inverse relationship between BF and BS 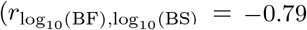 for TATA-box genes and − 0.82 for non-TATA-box genes), as well as positive and negative associations of GA with BS 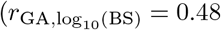 for TATA-box genes and − 0.64 for non-TATA-box genes) and with BF (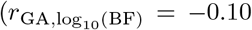 for TATA-box genes and −0.35 for non-TATA-box genes), respectively (Fig. 3K). These strong within-gene correlations differ from, and in some cases oppose, the trends observed in the posterior means (Fig. 3F), suggesting weak identifiability of burst kinetics.

Notably, the full posterior reveals differences in gene activity for non-TATA-box and TATA-box genes. Non-TATA-box genes reveal a bimodal GA distribution (Fig. 3K), suggesting that these genes occupy two distinct activity regimes, a feature that is obscured when considering only posterior means and also not observed for TATA-containing genes, which generally have lower GA (compare Fig. 3B). This suggests that promoter architecture may affect not only bursting behavior but also the range of accessible transcriptional states, with non-TATA-box promoters exhibiting greater heterogeneity in gene activity than TATA-containing promoters. Together, these results highlight the importance of gene activity in modulating burst kinetics.

### D. SFABC Quantifies Absolute Burst Kinetics using Time-resolved Transcriptomics

Next, we use SFABC for uncertainty quantification in the time-resolved analysis of metabolic labeling data^70^. We consider three models associated with different data modalities. The first is a steady-state telegraph model that uses only the total mRNA count (*u* + *l* modality), the second is a transient telegraph model that describes the dynamics of newly labeled mRNA alone (*l*), and the third is a metabolic labeling model that decomposes transcription into labeled and unlabeled mRNAs (*u* and *l*). See Fig. 4A and *SI Text 5*. We infer all four rate parameters *k*_on_, *k*_off_, *k*_syn_, *k*_deg_ and the labeling efficiency *λ* under the same priors (see caption of Fig. 4B).

**FIG. 4.**
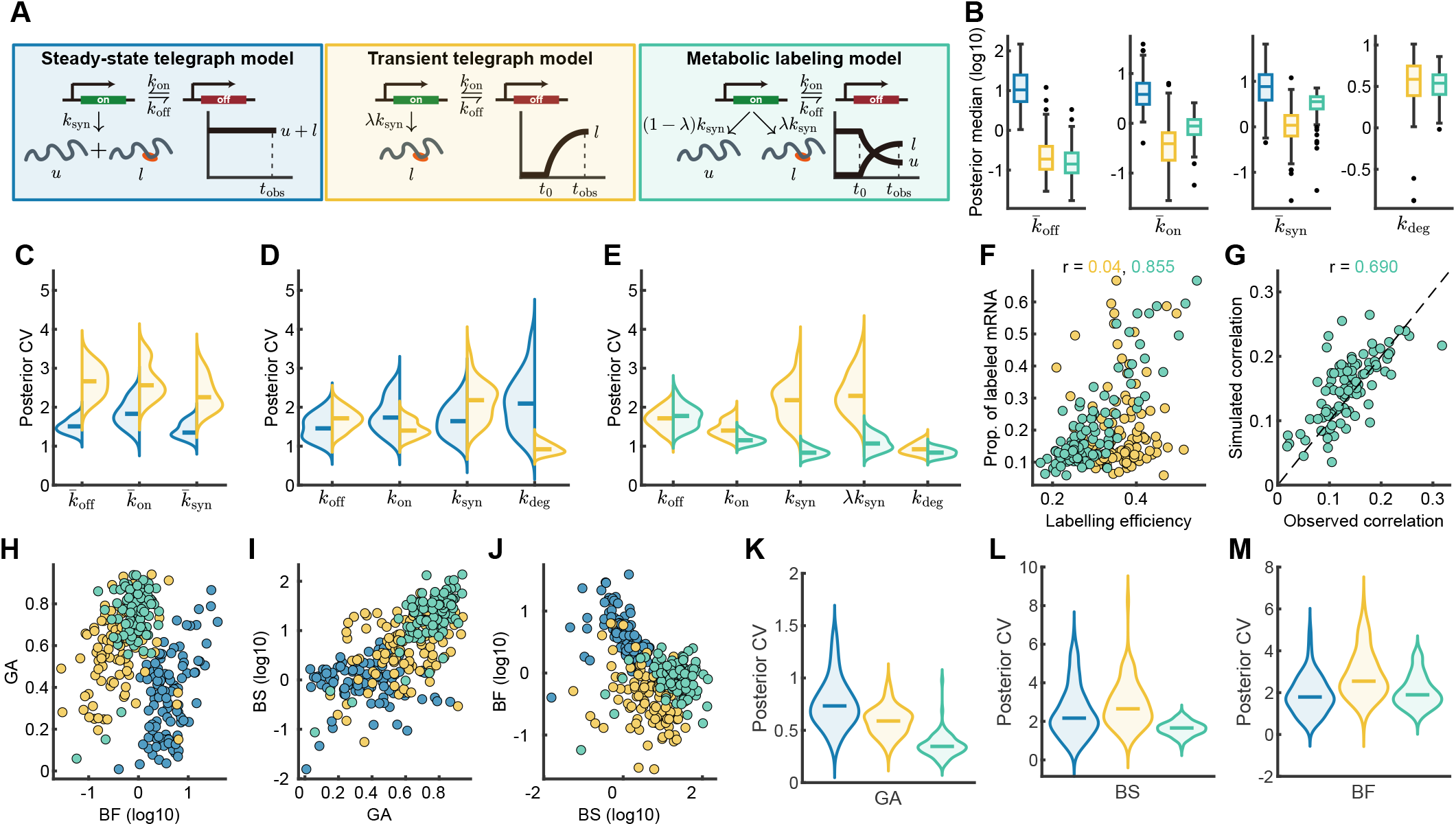
Inference of Burst Kinetics and Labeling Efficiency Using Time-resolved Transcriptomics. **(A)** Illustrations of stochastic reaction networks and data modalities. All models are fitted to primary mouse fibroblasts 4sU-labelled scRNA-seq data in Ref.^70^ that provides labeled and unlabeled counts *l* and *u* at time-point *t*_obs_ = 2*h*. The steady-state telegraph model considers their sum *u* + *l*, the transient telegraph uses only the labeled count *l*, and the metabolic labeling model involves paired observations (*u, l*). Degradation of labeled and unlabeled mRNA molecules occurs at a rate *k*_deg_ in all models. **(B)** Posterior median of the scaled switching rates 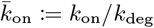 and 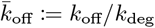, the scaled effective production rate 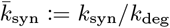 for steady-state telegraph and metabolic labeling models and 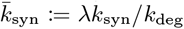 for transient telegraph model, and the degradation rate *k*_deg_ in the transient models. In all three models, we use priors *k*_off_, *k*_on_ ∼ 10^Uniform(*−*1.5,1.5)^ and *k*_syn_ ∼ 10^Uniform(*−*1.5,3)^. The prior of the labeling efficiency *λ* is 10^Uniform(*−*1,0)^ in the transient models. Inference is performed on a subset of 100 genes (gene selection details in *SI Text 6*), and inference setups are provided in S9 - S11 of *SI Table S1*. **(C)-(D)** Posterior CVs of scaled parameters and original parameters with steady-state and transient telegraph models. **(E)** Posterior CVs of original parameters with transient telegraph and metabolic labeling models. **(F)** Proportion of labeled mRNA in observed data, averaged over cells, versus the posterior mean of inferred labeling efficiency *λ* in transient telegraph and metabolic labeling models. The Pearson correlation coefficient (*r*) between them is reported for each model. **(G)** Correlation between labeled and unlabeled counts in posterior predictive data against that in observed data. **(H)-(J)** Pairwise plots of posterior median of burst kinetics GA, BF, and BS (calculated from the scaled parameters 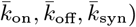. **(K)-(L)** Posterior CVs of burst kinetics GA, BF, and BS.

We adapt SFABC to the time-resolved setting by transforming the transient CME (3) into algebraic constraints that link the rate parameters with snapshot observations. To this end, we introduce an exponential time-weighting and integrate between time points *t*_1_ and *t*_2_ with *t*_2_ *> t*_1_ *≥* 0 as in^72^. The result is:

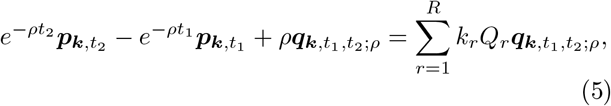

where 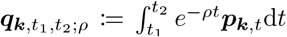 is not observable as it involves missing information between observation times. The transient CME assumes initial conditions with zero labeled transcripts (*l*) and a steady-state number of unlabeled transcripts (*u*). The latter obeys the steady-state CME (4), as does the total mRNA (*u* + *l*) at all times. The resulting relaxation can be efficiently solved using linear programming (see details in *SI Text 2*).

We used SFABC-SMC to infer the corresponding rate parameters of the three models and found that all models fit their respective data modalities (posterior predictive checks in ***SI Fig. S5A***). We found that the inference of posterior medians for relative parameters varied substantially across models and data modalities (Fig. 4B). Notably, the estimates from the metabolic labeling model tended to lie between those of the other two models, whereas degradation rates were consistent with those inferred from the transient telegraph model.

We found that parameter identifiability also varied substantially across models. Posterior uncertainty of the transient telegraph model was significantly worse than in the steady-state case (Fig. 4C). The telegraph model clearly identified degradation rates, but the identifiability of the other absolute rate parameters did not significantly improve over the steady-state telegraph model (Fig. 4D), even though the latter can only partially identify these absolute parameters due to its snapshot nature. The metabolic labeling model, which integrates labeled and unlabeled counts, yielded tighter posteriors for all parameters on an absolute timescale except *k*_on_, and thus provided more reliable inference (Fig. 4E). Improved identifiability in the metabolic model was due to incorporating labeling efficiency *λ*, which was predictive of the labeled mRNA fraction *λ* (Fig. 4F). Metabolic labeling model simulations also recovered the labeled-unlabeled correlations across genes (*r* = 0.69, Fig. 4G), validating that the model successfully integrates both transient and snapshot data modalities.

Data modality had a marked effect on inferred burst kinetics (Fig. 4H–J). Although burst size and burst frequency exhibited the expected inverse relationship, the steady-state and transient telegraph models inferred a broad range of gene activities and burst sizes, with median burst parameters strongly correlated across genes. By contrast, the metabolic-labeling model inferred uniformly higher gene activity with substantially reduced gene-to-gene variability. The identifiability gain was accompanied by positive correlations between degradation rate and both gene activity and burst size, which were less pronounced in the transient model (***SI Fig. S5D–F***). Consistent with these observations, the metabolic-labeling model substantially reduced posterior uncertainty in gene activity and burst size relative to the other models and data modalities (Fig. 4K–M). These results highlight that exploiting the joint distribution of time-resolved data significantly improves the identifiability of transcriptional burst kinetics.

### E. SFABC Identifies Cell Cycle-Dependent Transcriptional Regulation

Transcriptional kinetics often vary across the cell cycle. As these effects are often ignored in transcriptome-wide studies, such undulations can be mistakenly reported as biological noise. Identifying the transcriptional processes that are regulated by the cell cycle is crucial for this purpose^73–75^. To address this, we analyze scRNA-seq data with cell cycle reporters in^76^ and use SFABC to perform model selection among telegraph models with explicit cell division and cell cycle-dependent rates. We consider four cell cycle-dependent models: constant rates and models with varying *k*_on_, *k*_syn_ or *k*_deg_ (illustrations in Fig. 5A).

**FIG. 5.**
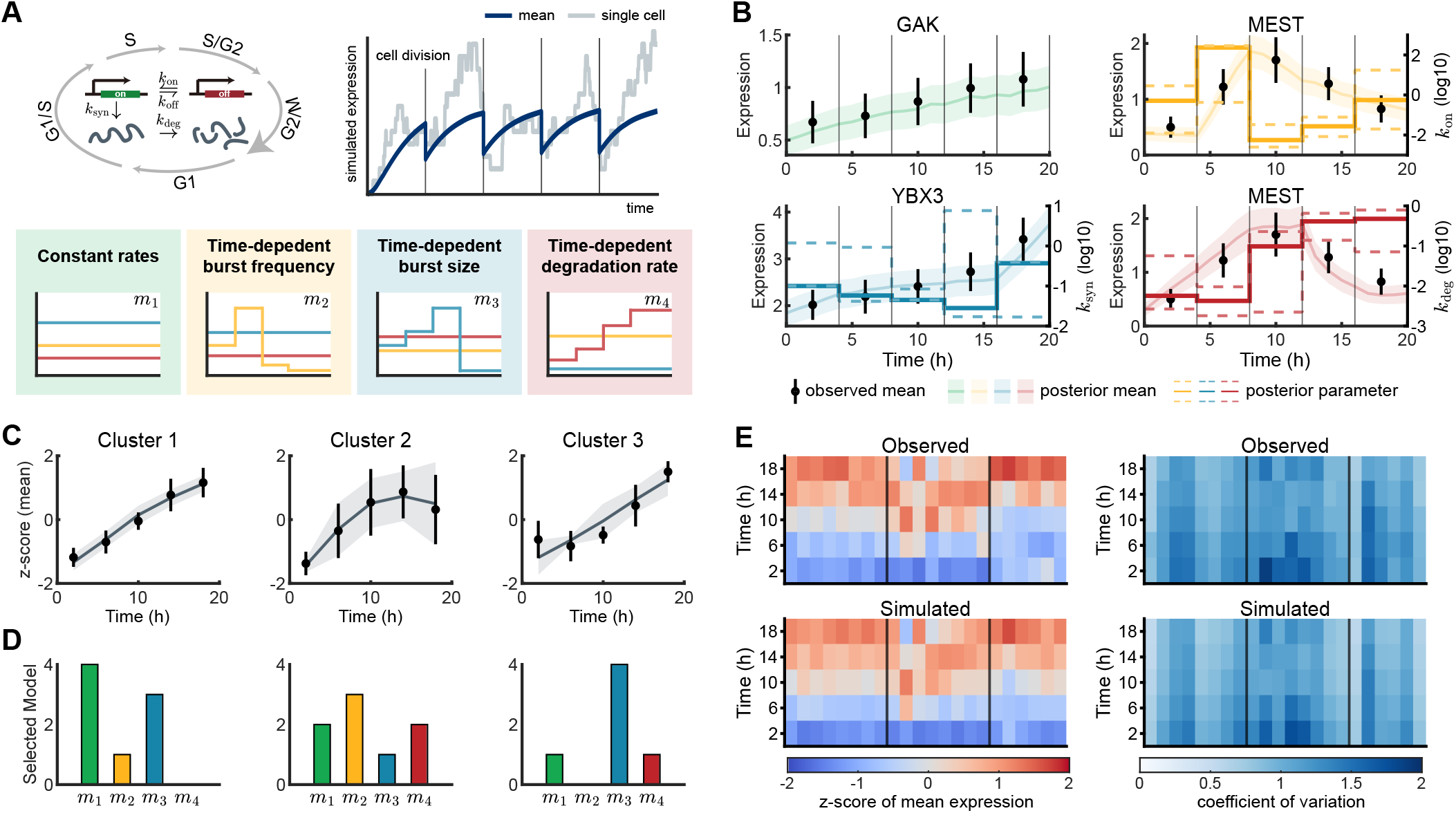
Model Selection among Telegraph Models with Cell Cycle-Dependent Rates. **(A)** Cartoon of cell cycle-dependent telegraph models. Model simulations indicate that expression approximately doubles over cell cycles on average, with significant fluctuations from cell to cell and over time. The model with constant rates *m*_1_ is the standard telegraph model, while *m*_2_, *m*_3_, and *m*_4_ are telegraph models with piecewise constant burst frequency (*k*_on_ varying), burst size (*k*_syn_ varying), and degradation rate (*k*_deg_ varying), respectively. **(B)** Representative gene-level fits and inferred time-dependent rate parameters to scRNA-seq data of RPE-1 cells with FUCCI cell cycle reporter (Ref.^76^; *SI Text 6*). We use assume parameters change discretely at timepoints {4, 8, 12, 16}, experimental observation taken at *T* = {2, 6, 10, 14, 18} and cell division occurs at *t*_div_ = 20 hours. For each example gene, black points show the observed mean mRNA counts at the sampled cell-cycle times, with error bars denoting the mean ± 0.2× standard deviation of the observed counts. Coloured solid curves show the posterior predictive mean expression from stochastic simulations of the selected telegraph model, and shaded bands show ± 0.2× the posterior predictive standard deviation. For models with a time-dependent parameter, the right y-axis shows the inferred log_10_ rate as a piecewise-constant function over the cell-cycle intervals; solid and dashed step curves denote the posterior median and the 95% credible interval, respectively. **(C)** Z-scores of the mean expression level in each cluster. Black points and error bars show the mean ±2 standard deviations across the observed gene profiles, while the grey curves and shaded regions show the mean ±2 standard deviations of the z-score obtained from the simulated posterior predictive. **(D)** Model selection summary for fitted genes in each cluster. For each gene, posterior model probabilities are computed from the accepted SFABC samples, and the selected model is defined as the model with the largest posterior probability (*SI Text 3*). Bars show the number of fitted genes assigned to each of the four candidate telegraph models within each cluster. **(E)** Observed and simulated z-scores of the mean expression level (left) and values of the coefficient of variation (right) per gene at the observation times. Vertical lines separate clusters. In (B) and (E), the simulation is run for 20 cell cycles to reach steady behavior before collecting posterior predictive samples for each parameter set drawn from the posterior. For each gene, 5,000 realizations from the posterior predictive distribution are simulated to plot the fitted expressions or to calculate the statistics. In all panels, we apply SFABC-SMC for model selection (*SI Text 3*).

To model the cell cycle progression, we use the transient equations (5) and assume parameters change discretely between observation times, the latter are akin to the phases (G1, G1/S, S, S/G2, G2/M) and cell division (M) at *t*_div_. We then impose

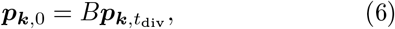

modeling equal partitioning of molecules at cell division.

The matrix 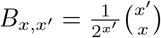 represents the probability for a cell to inherit *x* transcripts of the *x*′ transcripts of the mother cell^77,78^.

We applied SFABC-SMC for model selection to a number of representative genes with distinct temporal profiles (Figure 5B). To this end, we identified patterns by *k*-means clustering of the data (*SI Text 6*), showing steadily increasing, plateauing, and delayed transcription profiles (Fig. 5C, dots). SFABC recovered the mean expression levels for each cluster (lines), and its credible intervals recapitulated the gene distribution (Fig. 5C, shaded). Model selection revealed that the clusters correspond to different rate modulations, with the constant models dominating the steadily increasing transcripts (cluster 1), while plateaued and delayed expression were associated with cell-cycle-dependent burst frequency (cluster 2) and burst size modulation (cluster 3, Fig. 5D), respectively. The latter profiles (cluster 2 & 3) were also associated with constant rates or cell-cycle-dependent degradation in some genes, indicating substantial heterogeneity in burst kinetics modulation (***SI Fig. S6***).

Using the best models, SFABC’s predictive posterior distributions accurately recovered distinct cell-cycle patterns, and credible intervals recapitulated the data uncertainty (Figure 5B). In some cases (MEST), transcript profiles were equally well fit by different models with different rate modulations, indicating identifiability issues between cell cycle-dependent burst frequency and degradation. Overall, the posterior simulations reproduce the cluster-specific cell cycle patterns across genes (Fig. 5E).

The agreement indicates that the inferred models capture key temporal patterns in both mean expression and coefficient of variation across genes. Together, these results demonstrate that extending SFABC accurately captures biological noise and enables model comparison to unravel cell-cycle-dependent burst kinetics and transcriptional regulation.

## III. DISCUSSION

In this work, we propose SFABC, a simulation-free approximate Bayesian computation framework for stochastic reaction networks. SFABC leverages the Chemical Master Equation (CME), which links reaction-rate parameters to the underlying probability distributions of the stochastic reaction network. Rather than relying on stochastic simulations or analytical solutions, SFABC introduces a Bayesian sampling scheme that tests whether the CME associated with a given prior-parameter sample is compatible with Dirichlet-resampled data. These compatibility tests are formulated as convex optimization problems and can therefore be solved efficiently using linear programming.

We prove that the approximate posterior obtained by SFABC converges to the true Bayesian posterior in the limit of vanishing tolerance and increasingly refined relaxation approximations (see *Materials and Methods*). We further show that SFABC provides accurate posterior estimates from both snapshot and time-resolved transcriptomic data. Across synthetic benchmarks, SFABC achieves inference accuracy comparable to established ABC approaches while substantially reducing computational cost at large sample sizes, and remains robust across a wide range of parameter settings. When applied to steady-state scRNA-seq data, SFABC identifies systematic differences in inferred rate parameters associated with TATA-containing promoters and reveals pronounced within-gene parameter correlations, highlighting sloppiness and non-identifiability in burst kinetics. For time-resolved metabolic-labeling data, we find that identifiability is significantly improved by inferring absolute rate parameters and by exploiting correlation patterns across data modalities. Finally, we demonstrate the use of SFABC for model selection by identifying cell cycle-dependent transcriptional regulation.

In these applications, SFABC is able to incorporate model assumptions in a fundamentally different way from simulation-based approaches such as standard ABC. In the metabolic-labeling model, the total mRNA count is assumed to be at steady state. Similarly, the cell cycle-dependent telegraph model assumes cyclo-stationary behavior across cell divisions. Enforcing such stationary assumptions in simulation-based inference is computationally costly, since stochastic simulations must be run for sufficiently long times, or across sufficiently many cell cycles, to approach the required stationary behavior. Alternatively, dedicated simulators must be employed^79^. In contrast, within our simulation-free approach, these assumptions translate into directly checkable equations, thereby reducing the computational burden associated with biological constraints.

SFABC also differs conceptually from rejection-based ABC in its use of distance functions and summary statistics. In SFABC, the distance function and summary statistics are computed over observed states, which act as test functions distinguishing candidate probability distributions. As a result, questions of statistical sufficiency arise in a technically different way from simulation-based frameworks, where distances are typically computed between simulated and observed summary statistics. However, issues of model mispecification and model selection may carry over to SFABC^80,81^.

Another fundamental distinction from approximate-likelihood approaches lies in how approximations are introduced. Finite-state projection, moment approximations, and synthetic likelihoods can induce systematic parameter bias^27^. By contrast, the relaxation strategy employed by SFABC is constructed to contain the true parameters and probability solutions at all sample sizes, but does so at the cost of wider posteriors and therefore greater uncertainty. Hybrid approaches that combine SFABC with finite-state projection schemes could improve computational performance, but may introduce bias; the convergence properties of such methods would therefore need to be reassessed^82^.

Although SFABC provides an efficient and flexible framework with provable convergence guarantees, its computational efficiency has several limitations. Whereas the cost of simulation-based methods is primarily driven by repeated simulations and typically scales with factors such as sample size, the cost of SFABC is governed by the complexity of the underlying optimization problem. This complexity grows exponentially with the number of molecular species, limiting the scalability of SFABC to more complex networks. In addition, the convergence guarantees require enlargement of the state space in both steady-state and transient settings, as well as refinement of the time mesh in the transient setting. These refinements increase the number of optimization variables and introduce a practical tradeoff between accuracy and computational cost, alongside the usual tolerance-dependent trade-off. Adaptive state-space exploration, which remove low-probability states, could help alleviate this computational burden^83^.

These limitations suggest several directions for extending and improving SFABC. One natural extension is to replace the full CME with moment equations, so that the optimization problem scales with the number and order of moments rather than with the size of the state space. This could substantially improve scalability to higher-dimensional networks, albeit at the cost of sacrificing the convergence guarantees associated with Dirichlet sampling of the likelihood in the probability-based formulation (see *Materials and Methods*). Another promising direction is to reinterpret the feasibility test through the lens of control theory, with the aim of characterizing feasible parameter regions more directly and reducing the need for CME relaxation approximations^84–86^. It would also be valuable to translate algorithmic developments from ABC into the SFABC framework, including adaptive sequential Monte Carlo algorithms^52^ and regression adjustments for posterior correction^87^.

More broadly, SFABC reframes Bayesian inference for stochastic reaction networks by showing that repeated simulation is not always necessary as the governing equations themselves impose informative constraints on the data-generating distribution. By replacing simulated trajectories with optimization-based compatibility tests, SFABC turns model equations into tractable inferential objects and provides a principled way to quantify uncertainty without solving the CME or approximating the likelihood. This perspective opens a new route towards simulation-free inference in complex biological systems, where rich single-cell measurements are increasingly available but direct likelihood evaluation and large-scale simulation remain major bottlenecks. In this sense, SFABC presents a step toward a more general class of inference methods that exploit the structure of dynamical models directly. We expect that this paradigm will be useful for the rational design and reverse engineering of biological systems.

## IV. MATERIALS AND METHODS

### A. Practical Computation of the Acceptance Region Volume

SFABC weighs accepted samples by the inverse volume of the corresponding acceptance region. Due to independence between time points, the volume is 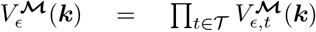 where 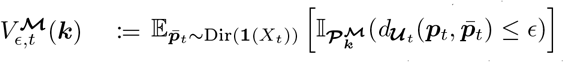 for *t* ∈ *T*. To estimate 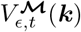 without repeated optimisation, we use a feasible solution of the CME relaxation 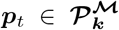, repeatedly sample 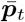 from the uniform Dirichlet distribution Dir(**1**(*X*_*t*_)), and take the proportion of samples with 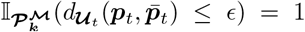 as our estimate, i.e., 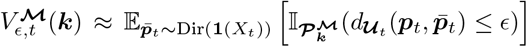, which does not affect convergence since ***p***_*t*_ → ***p***_***k***,*t*_ as ***M*** → ***X***.

### B. Convergence Proof of SFABC-Rej

The distribution of the accepted samples in SFABC-Rej is

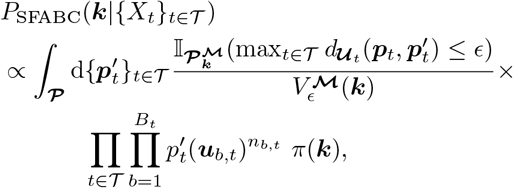

where ***P***:= ***P***_1_ *×* … … … *×* ***P***_*T*_ is the joint support of the Dirichlet distributions over observation times such that 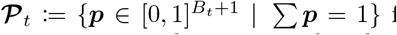 for *t*. To show convergence to the posterior distribution, we assume that the CME (3) admits a unique solution for every parameter vector ***k*** and we take the limits: (1) as ***M*** → ***X*** (and max_*i*_(*t*_*i*+1,opt_ *− t*_*i*,opt_) → 0 for transient data) the set 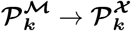 is a singleton *{****p***_***k***,*t*_*}*; and (2), we let the tolerance *ϵ* → 0. We then have

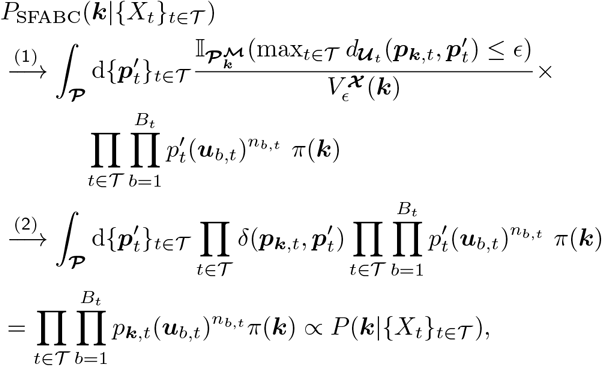

where the last line is the posterior in (2), which completes the proof.

## AUTHOR CONTRIBUTIONS

Author contributions: Z.L., M.B and P.T. designed the research, interpreted results and wrote the paper; Z.L. performed the derivations, designed the computations and analyzed the data.

## ACKNOWLEDGMENTS

ZL acknowledges support from the Roth Scholarship awarded by the Department of Mathematics at Imperial College London; MB is supported by EPSRC through the EPSRC Centre for Mathematics of Precision Healthcare (EP/N014529/1); PT is supported by UKRI through a Future Leaders Fellowship (MR/T018429/1 and MR/Y034309/1).

