## Supplementary Material for "Simulation-free approximate Bayesian computation for stochastic reaction networks"

Zekai Li,[1] Mauricio Barahona,[1, a] and Philipp Thomas[1, a]
[1]*Department of Mathematics, Imperial College London, London SW7 2AZ, United Kingdom*

(Dated: 19 October 2026)

### S1. BASELINE ABC ALGORITHMS

The ABC rejection algorithm (ABC-Rej)[1,2] works as follows to approximate the posterior:

**Algorithm S1: ABC rejection sampling (ABC-Rej)**

*S1* Sample a candidate parameter vector $\boldsymbol{k}$ from the prior $\pi(\cdot)$.
*S2* Simulate datasets $\{X'_t\}_{t\in\mathcal{T}}$ from the model with the parameter vector $\boldsymbol{k}$ at observation times $\mathcal{T}$.
*S3* For a given tolerance $\epsilon$, accept $\boldsymbol{k}$ if $\max_{t\in\mathcal{T}} d(X'_t, X_t) \leq \epsilon$ with a distance function $d$. Otherwise, go to *S1*.

ABC with sequential Monte Carlo (ABC-SMC) has been proposed to improve sampling efficiency by leveraging the ABC-Rej algorithm[3]. To collect $N$ samples from the posterior with a desired tolerance $\epsilon$, the SFABC-SMC algorithm samples a series of populations of weighted samples, $\boldsymbol{\mathcal{K}}_l := \{\boldsymbol{k}_l^{(i)}\}_{i=1}^N$ with weights $\boldsymbol{w}_l := \{w_l^{(i)}\}_{i=1}^N$ for $l = 1, \dots, L$. Each population contains samples accepted with shrinking tolerances $\epsilon_1 > \dots > \epsilon_L = \epsilon$, where $\{\epsilon_l\}_{l=1}^L$ is a chosen tolerance sequence. Given a tolerance sequence $\{\epsilon_l\}_{l=1}^L$ and perturbation kernels $\{\Psi_l\}_{l=1}^L$, the ABC-SMC algorithm proceeds as follows:

**Algorithm S2: ABC with sequential Monte Carlo sampling (ABC-SMC)**

*S1* Set tolerances $\epsilon_1, \dots, \epsilon_L$. Set population indicator $l = 1$.
*S2.0* Set sample count $i = 1$.
*S2.1* If $l = 1$, sample $\boldsymbol{k}$ from the prior $\pi(\cdot)$.
Else, sample $\boldsymbol{k}'$ from the previous population $\boldsymbol{\mathcal{K}}_{l-1}$ with weights $\boldsymbol{w}_{l-1}$. Perturb $\boldsymbol{k}'$ to obtain $\boldsymbol{k} \sim \Psi_l(\cdot|\boldsymbol{k}')$ where $\Psi_l$ is a perturbation kernel. If $\pi(\boldsymbol{k}) = 0$, return to *S2.1*.
*S2.2* Simulate datasets $\{X'_t\}_{t\in\mathcal{T}}$ from the model with the parameter vector $\boldsymbol{k}$ at observation times $\mathcal{T}$. For tolerance $\epsilon_l$, accept $\boldsymbol{k}$ if $\max_{t\in\mathcal{T}^*} d(X'_t, X_t) \leq \epsilon_l$ with a distance function $d$. Otherwise, go to *S2.1*.
*S2.3* If $i < N$, set $i = i + 1$, return to *S2.1*.
Else, calculate the weight of each sample $\boldsymbol{k}_l^{(i)}$:

$$w_l^{(i)} = \begin{cases} 1, & \text{if } l = 1 \\ \frac{\pi(\boldsymbol{k}_l^{(i)})}{\sum_{j=1}^N w_{l-1}^{(j)} \Psi_l\left(\boldsymbol{k}_l^{(i)} | \boldsymbol{k}_{l-1}^{(j)}\right)}, & \text{if } l > 1 \end{cases}$$

*S3* If $l < L$, set $l = l + 1$ and go to *S2.0*.

### S2. FEASIBILITY TEST IN SFABC

One of the core ideas of SFABC is the feasibility of the sampled parameter set and the sampled candidate state probability. In practice, this feasibility problem is often intractable because evaluating the state probability $\boldsymbol{p}_{\boldsymbol{k},t}$ requires solving the CME. This feasibility problem can be formulated as finding a feasible solution within the sets specified by the CME constraint and other constraints, but these sets may be infinite-dimensional due to an unbounded state space. Therefore, we relax the constraints in the set to obtain a solvable formulation.

The first approximation is to truncate the state space. We write the following CME over truncated state space $\mathcal{M} \subseteq \mathcal{X}$:

$$\frac{\mathrm{d}\boldsymbol{p}_{\boldsymbol{k},t}^{\tilde{\mathcal{M}}}}{\mathrm{d}t} = \sum_{r=1}^{R} k_r Q_r \boldsymbol{p}_{\boldsymbol{k},t}^{\mathcal{M}} \tag{S1}$$

for $\boldsymbol{p}_{\boldsymbol{k},t}^{\tilde{\mathcal{M}}} \coloneqq (P_t(\boldsymbol{x}|\boldsymbol{k}))_{\boldsymbol{x}\in\tilde{\mathcal{M}}}$ and $\boldsymbol{p}_{\boldsymbol{k},t} \coloneqq (P_t(\boldsymbol{x}|\boldsymbol{k}))_{\boldsymbol{x}\in\mathcal{M}}$ where $\tilde{\mathcal{M}}$ is the maximal subset of $\mathcal{M}$ which does not depend on states outside $\mathcal{M}$[4]. Here, the coefficient matrix $Q_r \in \mathcal{R}^{|\tilde{\mathcal{M}}|\times|\mathcal{M}|}$ has entries $Q_r(\boldsymbol{x},\boldsymbol{y}) = a_r(\boldsymbol{y})(\mathbb{I}_{\boldsymbol{y}+\boldsymbol{v}_r}(\boldsymbol{x}) - \mathbb{I}_{\boldsymbol{y}}(\boldsymbol{x}))$ for $\boldsymbol{x} \in \tilde{\mathcal{M}}$ and $\boldsymbol{y} \in \mathcal{M}$. In the rest of this section, we simplify the notation and write $\tilde{\boldsymbol{p}}_{\boldsymbol{k},t} \coloneqq \boldsymbol{p}_{\boldsymbol{k},t}^{\tilde{\mathcal{M}}}$ and $\boldsymbol{p}_{\boldsymbol{k},t} \coloneqq \boldsymbol{p}_{\boldsymbol{k},t}^{\mathcal{M}}$ for the truncated solutions to the CME with parameter $\boldsymbol{k}$, and denote by $\tilde{\boldsymbol{p}}_t$ and $\boldsymbol{p}_t$ a general probability vector over at time $t$ subsets $\tilde{\mathcal{M}}$ and $\mathcal{M}$, respectively. We assume the projection of $\mathcal{M}$ onto the observed species contains $\mathcal{U}_t$ for all $t \in \mathcal{T}$

At the steady state, the dynamics of the network no longer evolve in time, and we have

$$\frac{\mathrm{d}\tilde{\boldsymbol{p}}_{\boldsymbol{k}}}{\mathrm{d}t} = \sum_{r=1}^{R} k_r Q_r \boldsymbol{p}_{\boldsymbol{k}} = 0, \tag{S2}$$

where the time subscripts on the probabilities are omitted. We assess the compatibility of the rate parameters $\boldsymbol{k}$ with a sampled state probability $\boldsymbol{p}' \sim \mathrm{Dir}(n_1+1,\ldots,n_B+1,1)$ by checking whether a feasible solution $\boldsymbol{p}$ exists in the following set:

$$\mathcal{P}_{\boldsymbol{k}}^{\mathcal{M}} = \left\{ \boldsymbol{p} \in \mathbb{R}_+^{|\mathcal{M}|\times 1} \;\middle|\; \begin{array}{l} \sum_{\boldsymbol{x}\in\mathcal{M}} \boldsymbol{p}(\boldsymbol{x}) \leq 1 \\ \sum_{r=1}^{R} k_r Q_r \boldsymbol{p} = 0 \\ d_{\mathcal{U}}(\boldsymbol{p},\boldsymbol{p}') \leq \epsilon \end{array} \right\}, \tag{S3}$$

where $\mathcal{U}$ is the set of unique states in the observed data and the distance function $d_{\mathcal{U}}(\boldsymbol{p},\boldsymbol{p}')$ compares the probabilities over $\mathcal{U}$ (potentially only over observable species $\mathcal{S}$).

In contrast to the steady-state case with the exact dynamical constraint, the CME constraints involve integrals over the unobserved trajectories between observation times, so they cannot be enforced exactly from snapshots alone. To link the rate parameters with snapshot observations, we multiply an exponential test function $e^{-\rho t}$ to both sides of the CME for a given exponential weighting coefficient $\rho$ and integrate by parts from $t_1$ to $t_2$ with $t_2 > t_1 \geq 0$, which yields:

$$e^{-\rho t_2}\tilde{\boldsymbol{p}}_{\boldsymbol{k},t_2} - e^{-\rho t_1}\tilde{\boldsymbol{p}}_{\boldsymbol{k},t_1} + \rho\int_{t_1}^{t_2} e^{-\rho t}\tilde{\boldsymbol{p}}_{\boldsymbol{k},t}\mathrm{d}t = \sum_{r=1}^{R} k_r Q_r \int_{t_1}^{t_2} e^{-\rho t}\boldsymbol{p}_{\boldsymbol{k},t}\mathrm{d}t, \tag{S4}$$

for a given coefficient $\rho \in \mathbb{R}$. Defining $\boldsymbol{q}_{\boldsymbol{k},t_1,t_2;\rho} \coloneqq \int_{t_1}^{t_2} e^{-\rho t}\boldsymbol{p}_{\boldsymbol{k},t}\mathrm{d}t$ and $\tilde{\boldsymbol{q}}_{\boldsymbol{k},t_1,t_2;\rho} \coloneqq \int_{t_1}^{t_2} e^{-\rho t}\tilde{\boldsymbol{p}}_{\boldsymbol{k},t}\mathrm{d}t$, we can rewrite (S4) as

$$e^{-\rho t_2}\tilde{\boldsymbol{p}}_{\boldsymbol{k},t_2} - e^{-\rho t_1}\tilde{\boldsymbol{p}}_{\boldsymbol{k},t_1} + \rho\tilde{\boldsymbol{q}}_{\boldsymbol{k},t_1,t_2;\rho} = \sum_{r=1}^{R} k_r Q_r \boldsymbol{q}_{\boldsymbol{k},t_1,t_2;\rho}. \tag{S5}$$

With observations at time points $\mathcal{T}$, we consider time points $\mathcal{T}_{\mathrm{opt}} = \{t_1,\ldots,t_{T'}\}$ with $0 \leq t_1 < \ldots < t_{T'} = t_{\max}$ in the optimization, where $t_{\max}$ denotes the largest observation time and we assume $\mathcal{T} \subseteq \mathcal{T}_{\mathrm{opt}}$, and we denote $\boldsymbol{q}_{i;\rho} \coloneqq \int_{t_i}^{t_{i+1}} e^{-\rho t}\boldsymbol{p}_t\mathrm{d}t$ and $\tilde{\boldsymbol{q}}_{i;\rho} \coloneqq \int_{t_i}^{t_{i+1}} e^{-\rho t}\tilde{\boldsymbol{p}}_t\mathrm{d}t$ for $i = 1,\ldots,T'-1$. For a chosen set of exponential weighting coefficients $\rho \in \mathcal{R}$ in which we assume $\rho = 0$ is contained, we can write the constrained set for the transient case as:

$$\mathcal{P}_{\boldsymbol{k}}^{\boldsymbol{\mathcal{M}}} = \left\{ \begin{array}{l|ll} & \sum \boldsymbol{p}_t \leq 1, & t \in \mathcal{T}_{\text{opt}} \\ \boldsymbol{p}_t \in \mathbb{R}_+^{|\boldsymbol{\mathcal{M}}|\times 1} \text{ for } t \in \mathcal{T}_{\text{opt}} & \sum \boldsymbol{q}_{i;0} \leq t_{i+1} - t_i, & i = 1, \ldots, T-1 \\ \boldsymbol{q}_{i;\rho} \in \mathbb{R}_+^{|\boldsymbol{\mathcal{M}}|\times 1} & \sum \boldsymbol{q}_{i;\rho} \leq (e^{-\rho t_i} - e^{-\rho t_{i+1}})/\rho, & i = 1, \ldots, T-1; \rho \neq 0 \in \mathcal{R} \\ \text{for } i = 1, \ldots, T' - 1 & e^{-(\rho_1-\rho_2)t_{i+1}} \boldsymbol{q}_{i;\rho_2} \leq \boldsymbol{q}_{i;\rho_1} \leq e^{-(\rho_1-\rho_2)t_i} \boldsymbol{q}_{i;\rho_2}, & i = 1, \ldots, T-1; \rho_1 < \rho_2 \in \mathcal{R} \\ \text{and } \rho \in \mathcal{R} & e^{-\rho t_{i+1}} \tilde{\boldsymbol{p}}_{t_{i+1}} - e^{-\rho t_i} \tilde{\boldsymbol{p}}_{t_i} + \rho \tilde{\boldsymbol{q}}_{i;\rho} = \sum_{r=1}^R k_r Q_r \boldsymbol{q}_{i;\rho}, & i = 1, \ldots, T-1; \rho \in \mathcal{R} \\ & d_{\boldsymbol{\mathcal{U}}_t}(\boldsymbol{p}_t, \boldsymbol{p}'_t) \leq \epsilon, & t \in \mathcal{T} \end{array} \right\}, \tag{S6}$$

where the state probability $\boldsymbol{p}'_t$ is obtained as a Dirichlet resampling of the data at time $t \in \mathcal{T}$. Here, the first four constraints ensure that the probability and integral variables are valid and properly normalized, while the fifth one is the CME constraint and the last one is the distance constraint. Since the truncated transient CME is a finite-dimensional linear ODE in time, enforcing the CME-derived constraints only on the intervals of $\mathcal{T}_{\text{opt}}$ is an ordinary time discretization, and as the mesh size $\max_i(t_{i+1,\text{opt}} - t_{i,\text{opt}}) \to 0$ the discrete constraints converge to the continuous-time constraint.

Model-specific constraints can also be incorporated into the feasibility check, such as a steady-state constraint on the initial total-mRNA distribution in the metabolic-labeling model or a binomial partitioning constraint relating the mRNA-count distributions immediately before and after cell division. Additionally, (S5) also holds for piecewise constant rate parameters, as we can include all times at which any parameter value changes in the set $\mathcal{T}_{\text{opt}}$.

### S3. SFABC WITH SEQUENTIAL MONTE CARLO

To improve sampling efficiency, we propose SFABC with a sequential Monte Carlo (SFABC-SMC) sampling scheme. Given a tolerance sequence $\{\epsilon_l\}_{l=1}^L$ and perturbation kernels $\{\Psi_l\}_{l=1}^L$, the SFABC-SMC proceeds as follows:

**Algorithm S3: SFABC sequential Monte Carlo (SFABC-SMC)**

*S1* Set tolerances $\epsilon_1, \ldots, \epsilon_L$. Set population indicator $l = 1$.

*S2.0* Set sample count $i = 1$.

*S2.1* If $l = 1$, sample $\boldsymbol{k}$ from the prior $\pi(\cdot)$.
Else, sample $\boldsymbol{k}'$ from the previous population $\boldsymbol{\mathcal{K}}_{l-1}$ with weights $\boldsymbol{w}_{l-1}$. Perturb $\boldsymbol{k}'$ to obtain $\boldsymbol{k} \sim \Psi_l(\cdot|\boldsymbol{k}')$ where $\Psi_l$ is a perturbation kernel. If $\pi(\boldsymbol{k}) = 0$, return to *S2.1.*

*S2.2* For each observation time $t \in \mathcal{T}$, sample a candidate state probability $\boldsymbol{p}'_t \sim \text{Dir}(\boldsymbol{\alpha}(X_t))$.

*S2.3* Test if there exists a CME relaxation $\{\boldsymbol{p}_t\}_{t\in\mathcal{T}} \in \mathcal{P}_{\boldsymbol{k}}^{\boldsymbol{\mathcal{M}}}$ satisfies $\max_{t\in\mathcal{T}} d_{\boldsymbol{\mathcal{U}}_t}(\boldsymbol{p}_t, \boldsymbol{p}'_t) \leq \epsilon_l$. If such a solution exists, accept $\boldsymbol{k}$. Otherwise, go to *S2.1.*

*S2.4* If $i < N$, set $i = i + 1$, return to *S2.1.*
Else, calculate the volume $V_{\epsilon_l}(\boldsymbol{k}_l^{(i)})$ (see *Materials and Methods* in the main text). Calculate the weight of each sample $\boldsymbol{k}_l^{(i)}$:

$$w_l^{(i)} = \begin{cases} 1/V_{\epsilon_l}(\boldsymbol{k}_l^{(i)}), & \text{if } l = 1 \\ \dfrac{\pi(\boldsymbol{k}_l^{(i)})}{V_{\epsilon_l}(\boldsymbol{k}_l^{(i)}) \sum_{j=1}^N w_{l-1}^{(j)} \Psi_l\left(\boldsymbol{k}_l^{(i)} | \boldsymbol{k}_{l-1}^{(j)}\right)}, & \text{if } l > 1 \end{cases}$$

*S3* If $l < L$, set $l = l + 1$ and go to *S2.0.*

The weighted population $\boldsymbol{w}_L \boldsymbol{\mathcal{K}}_L$ can be shown to approximate the distribution of accepted samples from an SFABC-Rej algorithm with tolerance $\epsilon_L = \epsilon$ in the way that the ABC-SMC algorithm approximates the ABC-Rej algorithm in[3].

The SFABC-SMC algorithm for parameter inference can be easily extended to model selection problems. Consider candidate models indexed by $\{1, \ldots, M\}$ and the model-specific parameters $\boldsymbol{k}(m)$ for $m = 1, \ldots, M$, we use $\phi(\cdot)$ to denote the prior over these models and use $\pi_m(\cdot)$ to denote the prior of rate parameters given model $m$. Given tolerance sequence $\{\epsilon_l\}_{l=1}^L$, each population may contain samples from any model, namely $\boldsymbol{\mathcal{K}}_l := \{\boldsymbol{k}_l^{(i)}\}_{i=1}^N$ with models $\boldsymbol{m}_l := \{m_l^{(i)}\}_{i=1}^N$ and weights $\boldsymbol{w}_l := \{w_l^{(i)}\}_{i=1}^N$ for $l = 1, \ldots, L$, where each $m_l^{(i)} \in \{1, \ldots, M\}$. The SFABC-SMC model selection algorithm works as follows:

**Algorithm S4: SFABC-SMC for Model Selection**

*S1* Set tolerances $\epsilon_1, \ldots, \epsilon_L$. Set population indicator $l = 1$.
*S2.0* Set sample count $i = 1$.
*S2.1* Sample a model $m$ from $\phi(\cdot)$.
If $l = 1$, Sample $\boldsymbol{k}$ from the prior $\pi_m(\cdot)$.
Else, sample $\boldsymbol{k}'$ from the previous population $\boldsymbol{\mathcal{K}}_{l-1}^{m}$ with weights $\boldsymbol{w}_{l-1}^{m}$. Perturb $\boldsymbol{k}'$ to obtain $\boldsymbol{k} \sim \Psi_l^m(\cdot|\boldsymbol{k}'(m))$ where $\Psi_l^m$ is a perturbation kernel for model $m$. If $\pi_m(\boldsymbol{k}) = 0$, return to *S2.1*.
*S2.2* For each observation time $t \in \mathcal{T}$, sample a candidate state probability $\boldsymbol{p}_t' \sim \mathrm{Dir}(\boldsymbol{\alpha}(X_t))$.
*S2.3* Test if there exists a CME relaxation $\{\boldsymbol{p}_t\}_{t\in\mathcal{T}} \in \boldsymbol{\mathcal{P}}_{\boldsymbol{k}}^{\boldsymbol{\mathcal{M}}}$ satisfies $\max_{t\in\mathcal{T}} d_{\boldsymbol{\mathcal{U}}_t}(\boldsymbol{p}_t, \boldsymbol{p}_t') \leq \epsilon_l$. If such a solution exists, accept $\boldsymbol{k}$. Otherwise, go to *S2.1*.
*S2.4* If $i < N$, set $i = i + 1$, return to *S2.1*.
Else, calculate the volumes $V_{\epsilon_l}(\boldsymbol{k}_l^{(i)})$. Calculate the weight of each sample $\boldsymbol{k}_l^{(i)}$:

$$w_l^{(i)} = \begin{cases} 1/V_{\epsilon_l}(\boldsymbol{k}_l^{(i)}), & \text{if } l = 1 \\ \dfrac{\pi_m(\boldsymbol{k}_l^{(i)})}{V_{\epsilon_l}(\boldsymbol{k}_l^{(i)}) \sum_{j:m_{l-1}^{(j)}=m} w_{l-1}^{(j)} \Psi_l^m\left(\boldsymbol{k}_l^{(i)}|\boldsymbol{k}_{l-1}^{(j)}\right)}, & \text{if } l > 1 \end{cases}$$

*S3* If $l < L$, set $l = l + 1$ and go to *S2.0*.

With tolerance $\epsilon_L$, the posterior model probability is approximated as

$$P_{\text{SFABC-SMC}}(m|\{X_t\}_{t\in\mathcal{T}}) \approx \frac{1}{N}\sum_{i=1}^{N} \mathbb{I}(m_L^{(i)} = m), \tag{S7}$$

and the posterior distribution of parameters conditional on model $m$ is approximated by the weighted population $\{w_L^{(i)}\boldsymbol{k}_L^{(i)}\}_{i:m_L^{(i)}=m}$.

### S4. IMPLEMENTATION DETAILS OF INFERENCE ALGORITHMS

#### A. Inference Setup

We summarize the inference setup for the ABC and SFABC algorithms in Table S1, providing details on prior distribution, tolerance, type of constraints in the SFABC algorithms, and the choice of time points $\mathcal{T}$ and weightings $\mathcal{R}$ in the transient cases of SFABC.

#### B. Choice of SMC perturbation kernels

In setups S2 and S4, the perturbation kernel used in the SMC samplers is

$$\Psi_l(\cdot|\boldsymbol{k}') \sim \mathrm{Uniform}(k' - r_l, k' + r_l), \tag{S8}$$

independently for the three inferred parameters with $r_1 = 2.5$ and $r_2 = 1$. For setups S6 - S11, we use a perturbation kernel

$$\Psi(\cdot|\boldsymbol{k}') \sim k' \times \mathrm{Uniform}(0.75, 1.25) \tag{S9}$$

independently for each parameter. For setup S12, we use the following perturbation kernel:

$$\Psi_l(\cdot|\boldsymbol{k}') \sim 10^{\mathrm{Uniform}(\log_{10}(k')-r, \log_{10}(k')+r)} \tag{S10}$$

with $r_1 = 0.5$ and $r_2 = 0.25$, independently for each parameter in the current model.

### C. State space

One also needs to choose the states in the optimization problem, namely the set $\boldsymbol{\mathcal{M}}$. For the telegraph model, we have $\boldsymbol{\mathcal{M}} = \{0,1\} \times \{0,\ldots,2x_{\max}\}$, where $x_{\max}$ is the largest observation. For the metabolic labeling model, we have $\boldsymbol{\mathcal{M}} = \{0,1\} \times \{0,\ldots,2u_{\max}\} \times \{0,\ldots,2l_{\max}\}$, where $u_{\max}, l_{\max}$ are the largest observations of unlabeled and labeled mRNA counts, respectively. In the application to metabolic labeling data (S9 - S11 in Table S1), the genes in D2 are ranked by the size of their joint state space, quantified as $\max(u) \times \max(l)$, and the 100 genes with the smallest size are selected.

### D. Summary statistics and distance functions

We use standard moment-based distance functions. As the distances are calculated per time point, we describe the calculation for a single time point in this section and omit the time subscripts. For $\boldsymbol{x} = (x_1,\ldots,x_S)^\intercal$, we define $\boldsymbol{x}^{\boldsymbol{\alpha}} := \prod_{s=1}^S x_s^{\alpha_s}$ for $\boldsymbol{\alpha} := (\alpha_1,\ldots,\alpha_S)^\intercal \in \mathbb{N}^S$ and denote the $\boldsymbol{\alpha}$-th moment of dataset $X = (\boldsymbol{x}_1,\ldots,\boldsymbol{x}_N)$ to be $\mu^{\boldsymbol{\alpha}}(X) := \frac{1}{N}\sum_{i=1}^N \boldsymbol{x}_i^{\boldsymbol{\alpha}}$.

For ABC, we use

$$d(X',X) = \max_{|\boldsymbol{\alpha}|\le D} \left| \frac{\mu^{\boldsymbol{\alpha}}(X') - \mu^{\boldsymbol{\alpha}}(X)}{\hat{\mathrm{se}}(\mu^{\boldsymbol{\alpha}}(X))} \right|, \tag{S11}$$

where $|\boldsymbol{\alpha}| := \sum_{s=1}^S \alpha_s$ is the order of the moment and $\hat{\mathrm{se}}(\mu^{\boldsymbol{\alpha}}(X))$ is the bootstrap estimate of the standard deviation of the sample moment with 1000 bootstrap samples.

For SFABC, the candidate state probability is over the observed states with an additional bin representing the sum of probabilities over the unobserved states. Defining the moments over observed states $\mu^{\boldsymbol{\alpha}}_{\mathrm{obs}}(\boldsymbol{p}) := \sum_{b=1}^B p(\boldsymbol{u}_b)\boldsymbol{u}_b{}^{\boldsymbol{\alpha}}$, we consider the distance function:

$$d_{\boldsymbol{\mathcal{U}}}(\boldsymbol{p}^{\boldsymbol{\mathcal{M}}}_{\boldsymbol{k}}, \boldsymbol{p}') = \max_{|\boldsymbol{\alpha}|\le D} \left| \frac{\mu^{\boldsymbol{\alpha}}_{\mathrm{obs}}(\boldsymbol{p}^{\boldsymbol{\mathcal{M}}}_{\boldsymbol{k}}(\boldsymbol{\mathcal{U}})) - \mu^{\boldsymbol{\alpha}}_{\mathrm{obs}}(\boldsymbol{p}'(\boldsymbol{\mathcal{U}}))}{\hat{\mathrm{se}}(\mu^{\boldsymbol{\alpha}}_{\mathrm{obs}}(\boldsymbol{p}'(\boldsymbol{\mathcal{U}})))} \right|. \tag{S12}$$

where $\boldsymbol{p}(\boldsymbol{\mathcal{U}})$ denotes the probability over the observed states $\boldsymbol{\mathcal{U}}$, and $\hat{\mathrm{se}}(\mu^{\boldsymbol{\alpha}}_{\mathrm{obs}}(\boldsymbol{p}'(\boldsymbol{\mathcal{U}})))$ is the standard deviation of the $\boldsymbol{\alpha}$-th moment estimated by 1000 samples from the Dirichlet distribution for $|\boldsymbol{\alpha}| > 0$. A remarkable difference from ABC's distance function, in which $|\boldsymbol{\alpha}| = 0$ leads to a trivial comparison, is that the case $|\boldsymbol{\alpha}| = 0$ in SFABC compares the probability over the unobserved states, and it is necessary to include this in the distance function for an unbiased posterior estimate. In all numerical experiments, we set $\hat{\mathrm{se}}(\mu^{\boldsymbol{0}}_{\mathrm{obs}}(\boldsymbol{p}'(\boldsymbol{\mathcal{U}})))$ to be three standard deviations estimated from 1,000 samples from the Dirichlet distribution, where $\boldsymbol{0} := (0,\ldots,0)^\intercal \in \mathbb{N}^S$. We choose $D = 5$ for the telegraph model (S1 - S10, S12 in Table S1) and $D = 3$ for the metabolic labeling model (S11 in Table S1) so that the total number of moments is the same.

### E. Exact MCMC Sampler for Steady-State Telegraph Model

The steady-state probability of the telegraph model follows the Beta-Poisson distribution[5,6], which can be used to evaluate the likelihood. For benchmarking, we implement the Metropolis-within-Gibbs algorithm, which provides exact samples from the posterior, and compare them with ABC and SFABC samples (S1-S4 in Table S1). We run 10 independent chains for 55,000 iterations, discard the first 5,000 as burn-in, and keep every 1,000th draw thereafter (thinning). For each chain we initialize $k_{\mathrm{off}} \sim \mathrm{Gamma}(3,1)$, $k_{\mathrm{on}} \sim \mathrm{Gamma}(10, 0.75)$, and $k_{\mathrm{syn}} \sim \mathrm{Gamma}(5,1)$, and sample latent variables $p_j \sim \mathrm{Beta}(k_{\mathrm{on}}, k_{\mathrm{off}})$ for $j = 1,\ldots,N$. Each MCMC iteration updates the blocks $(\boldsymbol{p}, k_{\mathrm{on}}, k_{\mathrm{off}}, k_{\mathrm{syn}})$ sequentially using random-walk Metropolis steps. First, we propose $\boldsymbol{p}' = \boldsymbol{p} + \sigma_p \boldsymbol{\eta}$ with $\eta_j \sim \mathcal{N}(0,1)$ and only consider component-wise proposals that satisfy $0 < p'_j < 1$; valid components are accepted with the usual Metropolis rule based on the log-posterior difference. Next, we update $k_{\mathrm{on}}$, $k_{\mathrm{off}}$, and $k_{\mathrm{syn}}$ in turn with proposals $k'_m = k_m + \sigma_{k_m} z$ (with $z \sim \mathcal{N}(0,1)$), rejecting immediately if $k'_m \le 0$ and otherwise accepting according to the corresponding log-posterior ratio. We aggregate the thinned samples across chains to form posterior draws of $\boldsymbol{k} = (k_{\mathrm{off}}, k_{\mathrm{on}}, k_{\mathrm{syn}})$.

### S5. LIST OF STOCHASTIC REACTION NETWORKS

There are two stochastic reaction networks considered in this paper: the telegraph model and the metabolic labeling model. The telegraph model involves the following interactions:

$$G_{\text{on}} \underset{k_{\text{on}} g_{\text{off}}}{\overset{k_{\text{off}} g_{\text{on}}}{\rightleftharpoons}} G_{\text{off}}, \quad \varnothing \xrightarrow{k_{\text{syn}} g_{\text{on}}} X, \quad X \xrightarrow{k_{\text{deg}} x} \varnothing, \tag{S13}$$

where $G_{\text{on}}$ and $G_{\text{off}}$ denote the on and off states of a gene, and $X$ represents the mRNA count. We have $g_{\text{on}}, g_{\text{off}} \in \{0, 1\}$ with $g_{\text{on}} + g_{\text{off}} = 1$, and $x \in \mathbb{N}$. Here, $k_{\text{off}}$ and $k_{\text{on}}$ are the switching rates between the on and off states, $k_{\text{syn}}$ is the synthesis rate of mRNA molecules, and $k_{\text{deg}}$ is the degradation rate.

The metabolic labeling model, as an extension of the telegraph model, has the following interactions:

$$G_{\text{on}} \underset{k_{\text{on}} g_{\text{off}}}{\overset{k_{\text{off}} g_{\text{on}}}{\rightleftharpoons}} G_{\text{off}}, \quad \varnothing \xrightarrow{(1-\lambda) k_{\text{syn}} g_{\text{on}}} U, \quad \varnothing \xrightarrow{\lambda k_{\text{syn}} g_{\text{on}}} L, \quad U \xrightarrow{k_{\text{deg}} u} \varnothing, \quad L \xrightarrow{k_{\text{deg}} l} \varnothing, \tag{S14}$$

where $U$ is the count of the unlabeled mRNAs and $L$ is the count of the labeled ones. The additional parameter $\lambda$ is the labeling efficiency.

### S6. FILTERING SCRNA-SEQ DATA

For dataset D1 of mouse fibroblasts from[7], we considered only the CAST allele and obtained the processed UMI counts from the accompanying repository[?] . First, we filtered out genes with more than 1% missing counts. For the remaining genes, we filled missing counts with zero and further removed genes with a mean UMI count not exceeding 1 across cells. The remaining 979 genes were also observed in dataset D2 of mouse fibroblasts from[8]. After inference with SFABC-SMC using setups S7 and S8 in Table S1, we attempted to collect 100 posterior samples for each gene within the computational budget. Only genes for which 100 samples were obtained at the lowest tolerance were used for Fig. 3, leaving 764 genes for D1 and 865 genes for D2. We analyzed the total count (labeled and unlabeled) in Fig. 3. For Fig. 4, we used D2 and further selected 100 genes with the smallest product of the maximum UMIs across cells of the unlabeled ($u$) and 4sU-labeled ($l$) mRNAs, as the computational cost of SFABC scales with the size of the joint state space.

For the RPE-1 dataset with cell cycle reporters in[9], we used the 2,364 cells under the pulse metabolic labeling condition and considered the total mRNA count. We first filtered the 11,848 genes to remove those with mean expression below one at all cell-cycle phases. For the remaining 2,987 genes, z-scores across the cell cycle were computed and clustered into three groups using $k$-means. Ten genes per cluster were then randomly sampled, yielding 30 genes for downstream SFABC analysis (gene names listed in Table S2).

| Setup | Model | Prior | Sampling Algorithm | Tolerance | Constraints | Transient Optimization |
|---|---|---|---|---|---|---|
| S1 | Telegraph | $k_{\text{off}} \sim \text{Gamma}(3,1)$<br>$k_{\text{on}} \sim \text{Gamma}(10,0.75)$<br>$k_{\text{syn}} \sim \text{Gamma}(5,1)$ | ABC-Rej | 2 | - | - |
| S2 | | | ABC-SMC | $\{10,5,2\}$ | - | - |
| S3 | | | SFABC-Rej | 2 | Steady-state | - |
| S4 | | | SFABC-SMC | $\{10,5,2\}$ | Steady-state | - |
| S5 | Telegraph | $k_{\text{off}}, k_{\text{on}}, k_{\text{syn}} \sim 10^{\text{Unif}(-1,1)}$ | SFABC-SMC | $\{10,5,2\}$ | Steady-state | - |
| S6 | | | ABC-SMC | $\{10,5,2\}$ | - | - |
| S7 | Telegraph | $k_{\text{off}}, k_{\text{on}} \sim 10^{\text{Unif}(-1.5,1.5)}$<br>$k_{\text{syn}} \sim 10^{\text{Unif}(-1.5,3)}$ | SFABC-SMC | $\{10,6,4\}$ | Steady-state | - |
| S8 | | | SFABC-SMC | $\{50,20,10\}$ | Steady-state | - |
| S9 | Telegraph | $k_{\text{off}}, k_{\text{on}}, k_{\text{deg}} \sim 10^{\text{Unif}(-1.5,1.5)}$<br>$k_{\text{syn}} \sim 10^{\text{Unif}(-1.5,3)}$ | SFABC-SMC | $\{50,20,10\}$ | Steady-state | - |
| S10 | Telegraph | $k_{\text{off}}, k_{\text{on}}, k_{\text{deg}} \sim 10^{\text{Unif}(-1.5,1.5)}$<br>$\lambda k_{\text{syn}} \sim 10^{\text{Unif}(-1.5,3)} \cdot 10^{\text{Unif}(-1,0)}$ | SFABC-SMC | $\{50,20,10\}$ | Transient | $\mathcal{T} = \{0,1,2\}$<br>$\lvert\mathcal{R}\rvert = 4$ |
| S11 | Metabolic labeling | $k_{\text{off}}, k_{\text{on}}, k_{\text{deg}} \sim 10^{\text{Unif}(-1.5,1.5)}$<br>$k_{\text{syn}} \sim 10^{\text{Unif}(-1.5,3)}$<br>$\lambda \sim 10^{\text{Unif}(-1,0)}$ | SFABC-SMC | $\{50,20,10\}$ | Steady-state† & transient | $\mathcal{T} = \{0,1,2\}$<br>$\lvert\mathcal{R}\rvert = 6$ |
| S12 | Cell-cycle telegraph | $k_{\text{off}}, k_{\text{on}} \sim 10^{\text{Unif}(-3,3)}$<br>$k_{\text{syn}} \sim 10^{\text{Unif}(-2,2)}$<br>$k_{\text{deg}} \sim 10^{\text{Unif}(-3,1.5)}$ | SFABC-SMC | $\{10,7.5,5\}$ | Transient & cell-cycle†† | $\mathcal{T} = \{0,0.5,1,\ldots,20\}$<br>$\lvert\mathcal{R}\rvert = 4$ |

TABLE S1. **Summary of Inference Setup.** For S1 - S8, we fix $k_{\text{deg}} \equiv 1$. † The total mRNA count (i.e., labeled + unlabeled) is required to follow a steady-state distribution at $t = 0$h as the initial condition. †† The cell-cycle telegraph model has a specific constraint forcing the distribution of mRNA count at $t = 0$h to follow a Binomial down-sampling with probability parameter 0.5 from that at $t = 20$h.

| **Cluster 1** | **Cluster 2** | **Cluster 3** |
|---|---|---|
| CENPB | VEGFB | KPNA2 |
| LTA4H | **MEST** | SPC25 |
| **GAK** | MYCBP2 | CENPF |
| PDZD11 | ADAMTSL1 | **YBX3** |
| PLEKHA5 | FBN1 | EVA1A |
| TXNL4A | CHAF1A | ARHGEF2 |
| FH | CTPS1 | PSMC3 |
| RPL4 | ARHGAP10 | RASSF3 |
| PABPC1 | SLC2A4RG | ALYREF |
| RPL5P17 | SSH1 | CDCA8 |

TABLE S2. **Genes Used for SFABC Analysis in the Cell-Cycle Dataset[9].** Related to Fig. 5. Genes were first filtered by mean expression, clustered into three groups based on their z-scored expression profiles across cell-cycle phases, and then ten genes were sampled from each cluster. Example genes in Fig. 5B are highlighted.

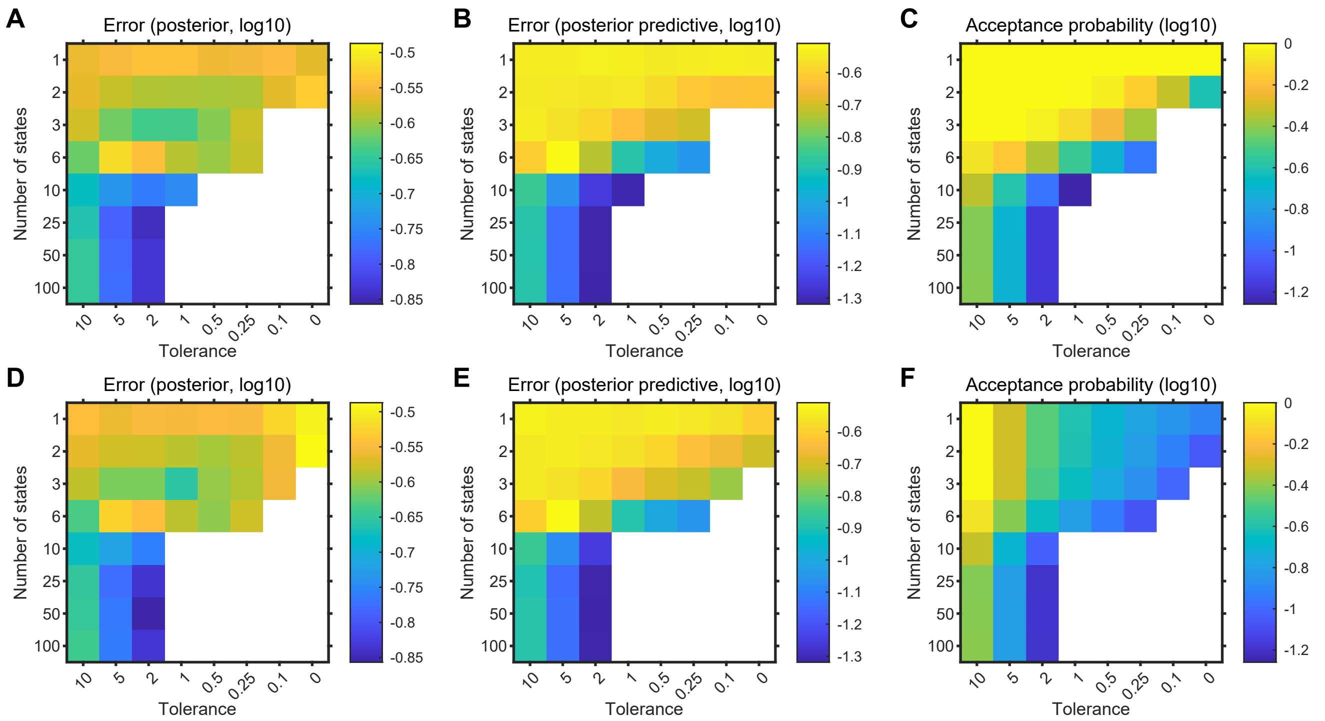


FIG. S1. **Effect of Truncated State-Space Size and Tolerance on SFABC Performance for Steady-State Telegraph-Model Data.** Heatmaps show validation metrics for the 20 synthetic datasets with sample size 500 used in Fig. 2B,C. The upper row shows SFABC-Rej (A–C), and the lower row shows SFABC-SMC (D–F). Columns show the posterior error relative to MCMC, the posterior predictive error relative to the observed data, and the acceptance probability, all on the $\log_{10}$ scale. Results are averaged over the 20 datasets for different tolerances and numbers of states included in the feasibility test. Empty cells indicate settings for which the required number of posterior samples was not obtained under the fixed optimization budgets: 15,000 optimizations for each SFABC-Rej run, and 25,000 optimizations for each SFABC-SMC run. While we expect the acceptance probability to decrease both as the tolerance is tightened and as the truncated state space is enlarged, small nonmonotonic fluctuations can occur because independent SFABC-Rej runs across different tolerances and truncation sizes used different random seeds. Except for the tolerance settings and truncated state-space sizes, the inference setup is the same as in S3-S4 in Table S1. In SMC, we use the perturbation kernel in (S8) with linearly decreased kernel parameter $r = \{2.5, 2.125, 1.75, 1.375, 1, 0.625, 0.25\}$.

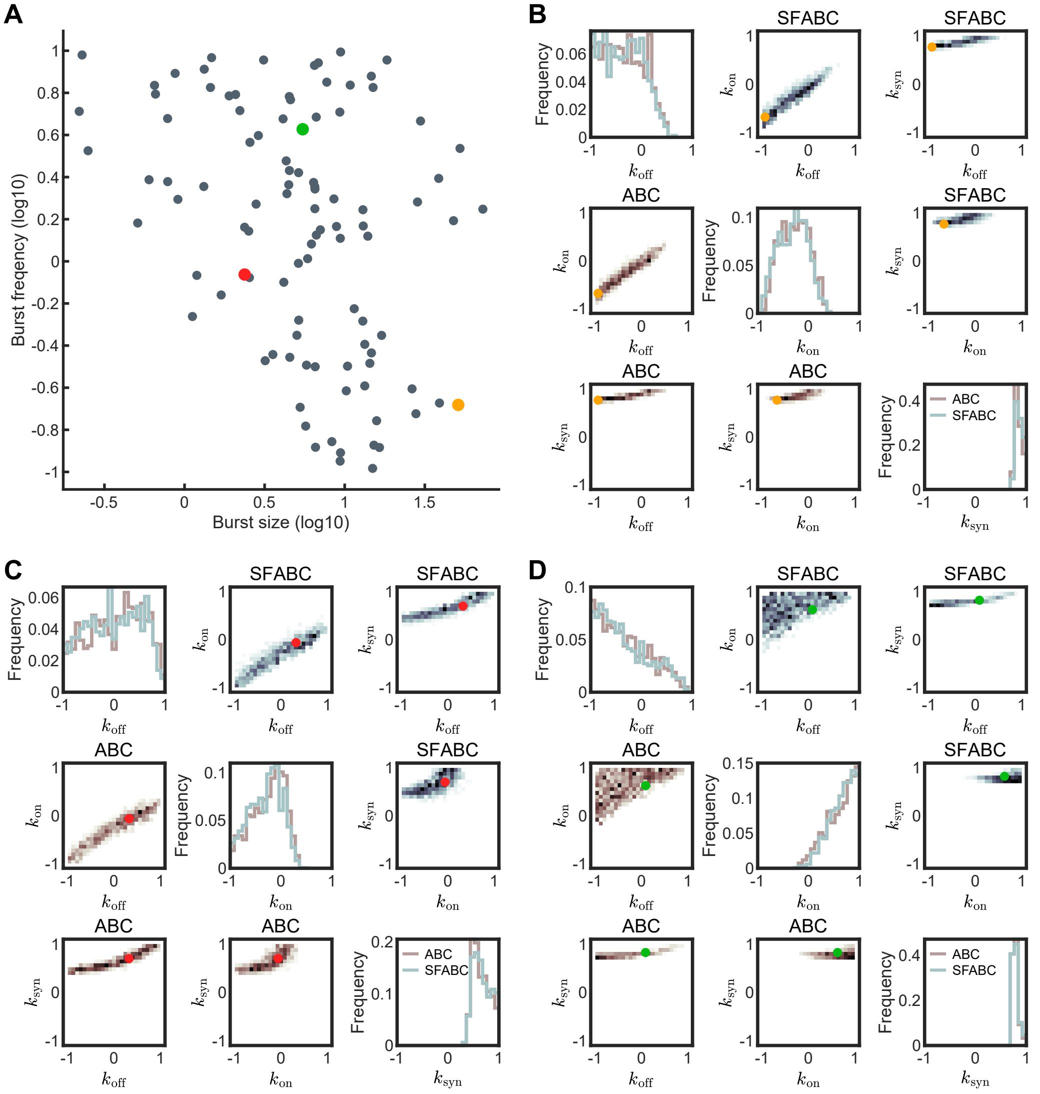


FIG. S2. **Comparison of ABC and SFABC Posterior Estimates for different synthetic data sets.** **(A)** Burst frequency and burst size of the 100 parameter sets used in the robustness test of SFABC in Fig 2E-H. Colored points represent the three example parameter sets in (B)-(D). **(B)-(D)** Posterior comparison between ABC and SFABC estimates. Diagonal sub-panels show marginal posterior distributions of the three parameters, and the other sub-panels compare the pairwise posterior distributions between them. In B-D, distributions are shown on a $\log_{10}$ scale.

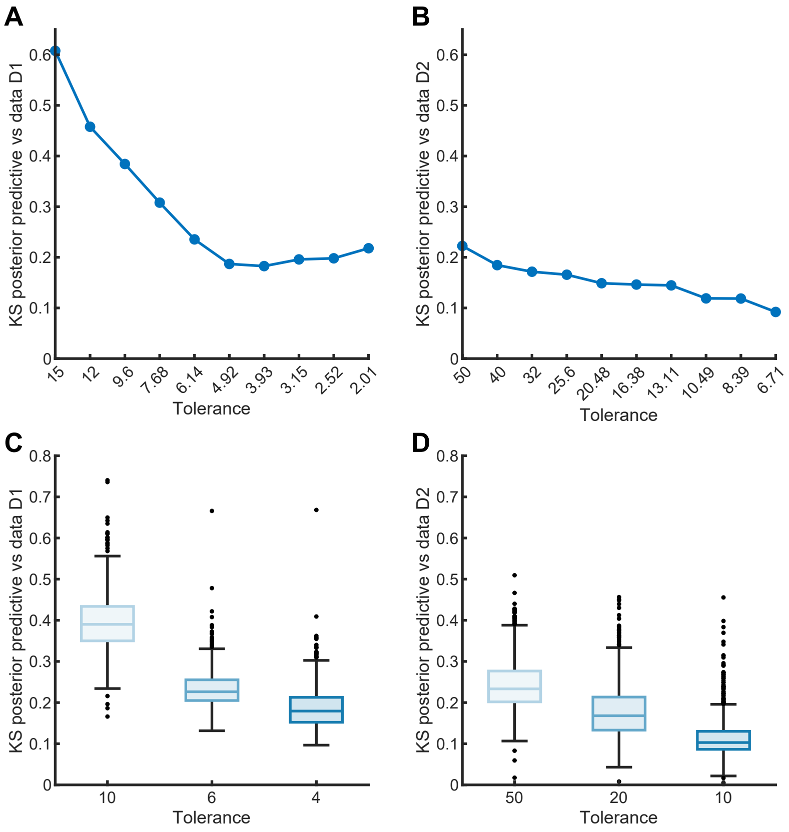


FIG. S3. **Selection of Tolerances and Validation of Posterior Estimates for Inference of the Telegraph Model with Steady-State scRNA-seq Datasets. (A)-(B)** Tolerance selection based on the Kolmogorov-Smirnov (KS) metrics. We test across ten tolerances for each dataset with SFABC-SMC, $\{15, 15 \times 0.8^1, \ldots, 15 \times 0.8^9\}$ for D1 and $\{50, 50 \times 0.8^1, \ldots, 50 \times 0.8^9\}$ for D2. The tolerance sequences selected for analysis in the main text are $\{10, 6, 4\}$ for D1 and $\{50, 20, 10\}$ for D2, balancing accuracy and computational cost. **(C)-(D)** KS metrics between SFABC's posterior predictives and observed datasets.

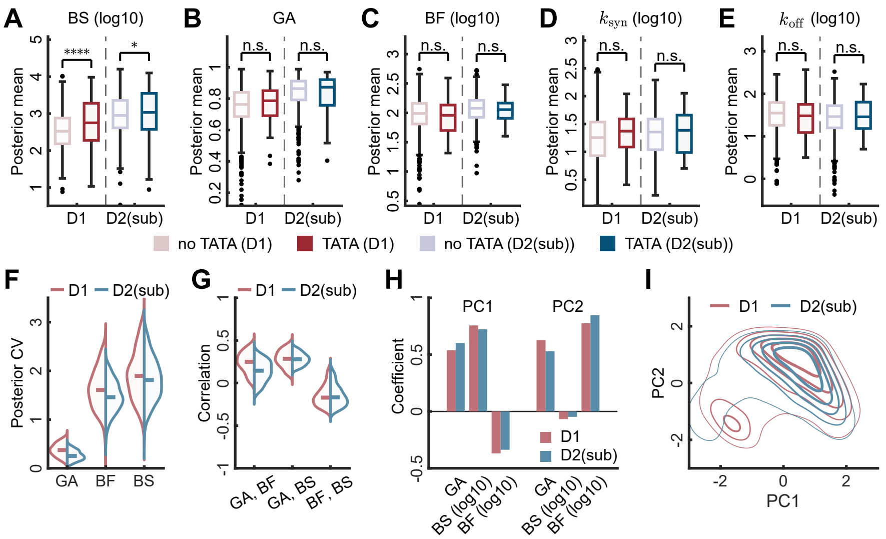


FIG. S4. **Ablation Study on the Larger Steady-State scRNA-seq Datasets.** Here, we randomly subsample the dataset D2 in[8] with 8,916 cells to obtain D2(sub) with 224 cells, the same as in D1 in[7]. **(A)-(E)** Posterior means of burst kinetics and individual rate parameters, grouped by dataset and gene type (with or without a TATA element). **(F)-(G)** Coefficients of variation (CVs) of marginal posterior distributions and pairwise correlations in posterior samples across genes in each dataset, restricted to the 10th-90th percentiles. **(H)** Coefficients of principle components. For each gene, principal component analysis is performed on the (across-gene) posterior samples of GA, $\log_{10}$(BS), and $\log_{10}$(BF) for each of D1 and D2. **(I)** Distribution of posterior samples projected to the plane of the first principal component (PC1) and the second one (PC2).

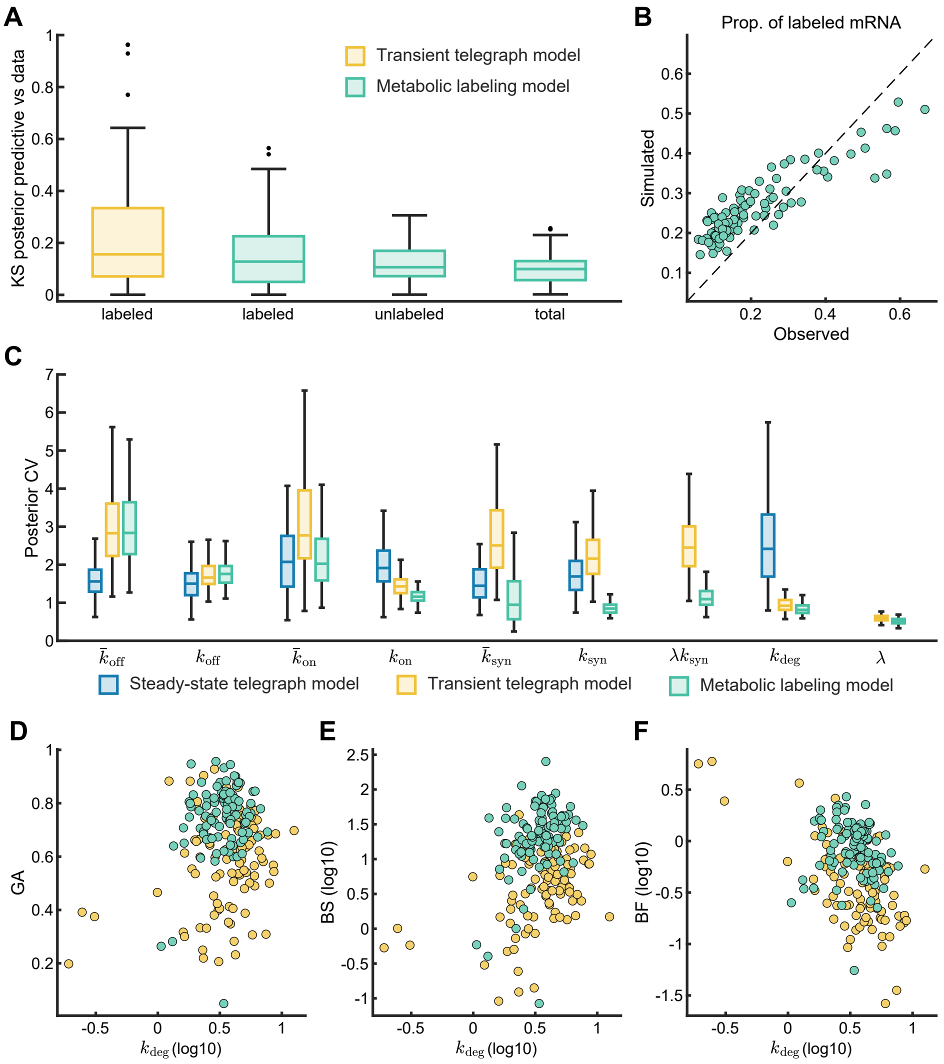


FIG. S5. **Validation of Posterior Estimates for Inference of the Transient Telegraph and Metabolic Labeling Models. (A)-(B)** KS metrics between the posterior predictive and the observed data. We compare the labeled counts for the transient telegraph model, and compare labeled, unlabeled, and total counts for the metabolic labeling model. **(B)** Proportion of labeled mRNA in total mRNA in observed and simulated data. **(C)** Posterior CVs for inferred kinetic parameters across the 100 genes. The scaled parameters are defined as $\bar{k}_{\text{on}} := k_{\text{on}}/k_{\text{deg}}$ and $\bar{k}_{\text{off}} := k_{\text{off}}/k_{\text{deg}}$, with $\bar{k}_{\text{syn}} := k_{\text{syn}}/k_{\text{deg}}$ for the steady-state telegraph and metabolic labeling models and $\bar{k}_{\text{syn}} := \lambda k_{\text{syn}}/k_{\text{deg}}$ for the transient telegraph model. Outlier markers are omitted for visual clarity. **(D)-(F)** Posterior median of burst kinetics gene activity (GA), burst size (BS), and burst frequency (BF) against that of the degradation rate.

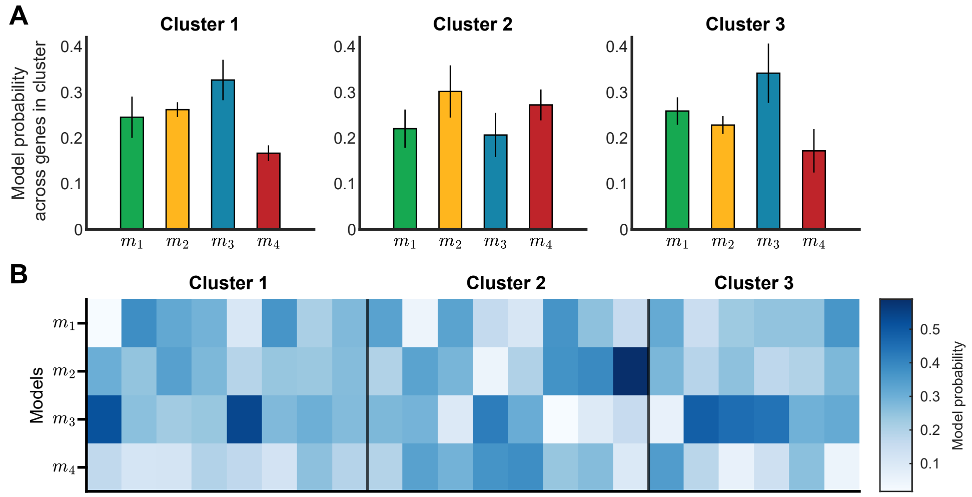


FIG. S6. **Model Probabilities Inferred for the Telegraph Model with Cell Cycle-Dependent Rates. (A)** Model probabilities across genes in each cluster (clusters defined as described in the main text). Bars show cluster-level mean model probabilities ± standard error of the mean across fitted genes. **(B)** Model probability per fitted gene.
